# The ancient metazoan cytoplasmic intermediate filament protein, Cilin, shapes cilia arrangement and tissue architecture

**DOI:** 10.1101/2025.08.21.671432

**Authors:** Stanislav Kremnyov, Nadezhda Rimskaya-Korsakova, Tatiana Lebedeva, Alena Kizenko, Lars Hering, Georg Mayer, Carl-Philipp Heisenberg, Andreas Hejnol

**Affiliations:** Institute of Zoology and Evolutionary Research, Faculty of Biological Sciences, Friedrich Schiller University Jena, Jena, Germany; Institute of Science and Technology Austria, Klosterneuburg, Austria; Department of Zoology, Institute of Biology, University of Kassel, Kassel, Germany

**Keywords:** Intermediate filaments, lamin, cilia, Ctenophora, *Mnemiopsis leidyi*

## Abstract

The emergence of animal multicellularity demanded novel cytoskeletal systems to support cellular architecture and tissue integrity. Cytoplasmic intermediate filaments (cIFs) are essential components of this scaffold, yet their evolutionary origins remain obscure. Here, we identify and characterize a lamin-derived, *bona fide* cIF protein in a ctenophore—a sister lineage to all other extant animals. We name this protein **Cilin**. Unlike nuclear lamins, Cilin lacks a nuclear localization signal and instead localizes predominantly to motile ciliary structures, including comb plates, the aboral organ, and sperm flagella, indicating a central role in ciliary architecture and function. Remarkably, Cilin is also present in non-ciliated cells, suggesting early functional diversification of cIFs in animal evolution. Phylogenetic and structural analyses position Cilin within an ancestral class of intermediate filament proteins, homologous to cnidarian and bilaterian nematocilins and lamin-tail-domain-containing (LMNTD) proteins. In humans, LMNTD proteins are enriched in ciliated epithelia and spermatids, pointing to a deeply conserved role in cilia-associated functions. These findings establish cilins as the earliest lamin-derived cytoplasmic intermediate filament proteins in metazoans, likely contributing to both ciliary function and the emergence of multicellular tissue organization.

## Introduction

The cytoskeleton of animals (Metazoa) is composed of three major filament systems— actin filaments, microtubules, and intermediate filaments (IFs)—which together orchestrate cellular shape, stability, and organization^1^. Unlike actin and microtubules, which are polar and require accessory proteins for dynamic assembly, IFs are non-polar polymers that self-assemble into flexible, mechanically robust networks^2^. IFs are essential for maintaining cellular and nuclear integrity, resisting mechanical stress, and anchoring organelles in the cytoplasm^3^. They can also act as scaffolds for signaling pathways involved in differentiation and development^4^.

All IF proteins share a conserved central α-helical rod domain, flanked by variable N-terminal (head) and C-terminal (tail) domains^5,6^. The rod domain of IFs is highly conserved and plays a central role in their structural assembly^6^. Among the large family of IF proteins, nuclear lamins date back to the last common ancestor of eukaryotes (plants, fungi and Metazoa)^7^, and are localized exclusively to the nucleus and are characterized by a nuclear localization signal (NLS), a C-terminal CaaX motif for inner nuclear membrane attachment, and an Ig-like lamin tail domain (LTD)^8,9^ (Fig. 1A). In contrast, all cytoplasmic IF (cIF) proteins lack NLS signal sequences and CaaX motifs and play diverse roles in the cytoplasm. An LTD domain may or may not be present^10^. Invertebrate cIF proteins often have a C-terminal domain that structurally resembles the LTD of lamins, including an Ig-fold^11^. In vertebrates, the Ig-fold is absent in all types of cIF proteins and is present only in lamins^11^ (Fig. 1A). Notably, in mammals, an LTD domain is also found in lamin-tail-domain-containing (LMNTD) proteins, which contain only this domain and lack the central rod domain entirely^12,13^. However, the phylogenetic relationship between LMNTD proteins and intermediate filaments remains unclear, as does their biological function.

**Figure 1.**
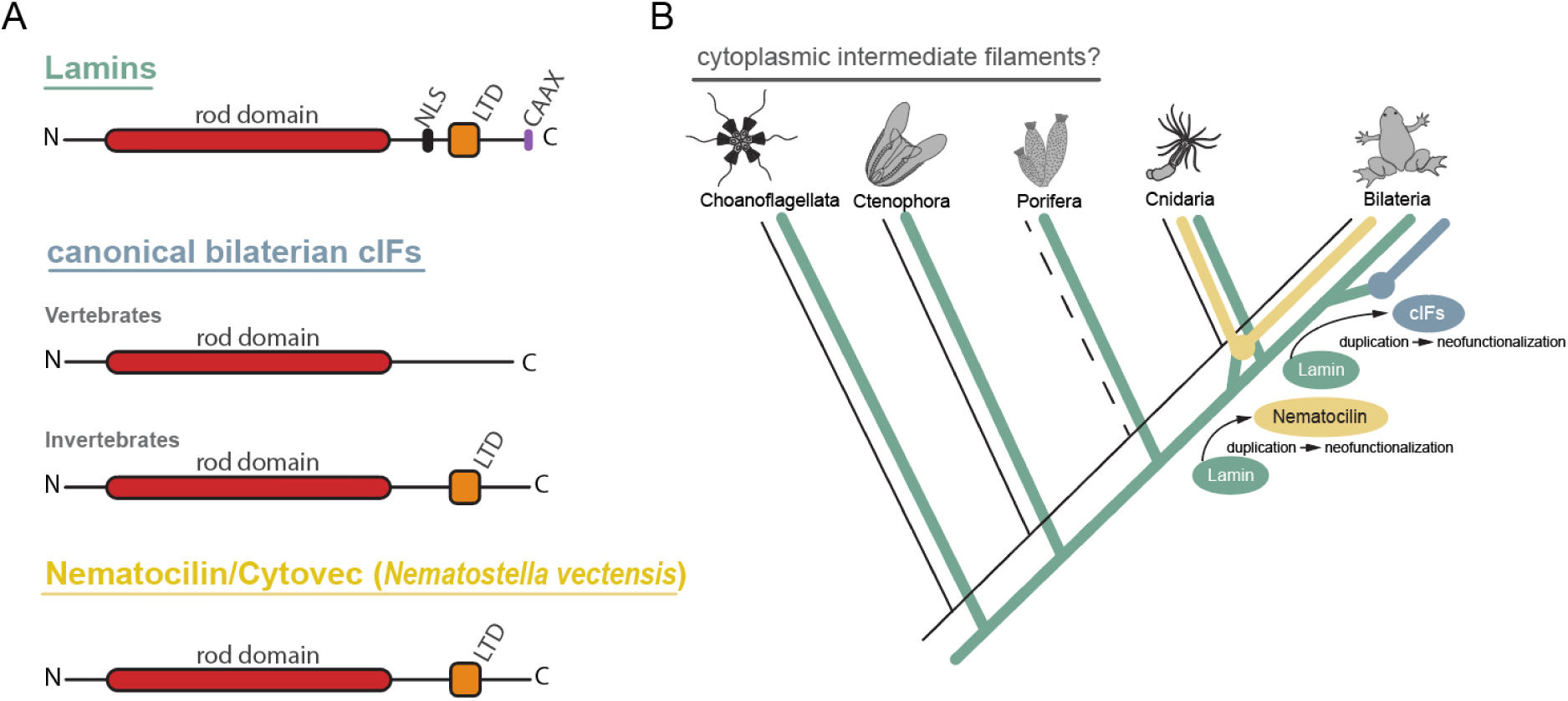
IF domain organization (A) and current scenario on the origin and evolution of cytoplasmic IF proteins in Metazoa (B). According to the current state of knowledge, the earliest appearance of cIFs in metazoans was the emergence of nematocilin in cnidarians (yellow), resulting from a gene duplication and subsequent neofunctionalization. The greatest diversity of cIFs arose in bilaterians (blue) through an independent duplication of lamin, followed by its neofunctionalization. CaaX - signal which targets proteins to membranes, LTD - Lamin Tail Domain, NLS - Nuclear Localisation Signals

While the evolutionary origins of actin and tubulin are well understood and likely trace back to prokaryotic ancestors^14–17^, the evolutionary history of IF proteins remains uncertain^18^. Lamins are thought to have emerged early in eukaryotes^19,7,8^, while cIF proteins are thought to have emerged later in multicellular animals through duplication and neofunctionalization of ancestral lamin genes^20,21^.

In bilaterians, cIFs are widely represented, mammals alone express over 70 different cIF genes^22^. However, they have been lost in panarthropods^23^ (tardigrades, onychophorans, and arthropods) and re-evolved from lamins in tardigrades^24^ and collembolans^25^. Among non-bilaterian animals, only a single type of cIF proteins has been identified, comprising nematocilin in hydrozoans^26^ and cytovec in anthozoans^27^ (Fig. 1A). Nematocilin was discovered in *Hydra magnipapillata* nematocytes, where it localizes specifically to the central filament of the cnidocil, a cilium-derived mechanosensory structure involved in triggering nematocyst discharge^26^. When first discovered, nematocilin was identified as not belonging to the group of canonical bilaterian cIFs^26^. However, later phylogenetic analyses showed that nematocilin-like genes are present in most bilaterians (except Ecdysozoa and Platyhelminthes)^7^, suggesting its shared evolutionary origin in the common ancestor of Cnidaria and Bilateria. The function of nematocilin-like proteins in bilaterians has yet to be determined. These proteins are thought to have evolved independently from lamins via gene duplication in the common ancestor of cnidarians and bilaterians, and are not homologous to the canonical cIFs present in bilaterians^7^ (Fig. 1B). Notably, no studies have investigated cIFs in other non-bilaterian animals, such as sponges or ctenophores, or in the closest metazoan relatives, the choanoflagellates. Therefore, it remains unclear whether the first cIFs evolved in the last common ancestor of cnidarians and bilaterians, or even earlier.

To resolve the ambiguity surrounding the origin of cIFs, we investigated intermediate filament proteins in ctenophores, the sister group to all other animals^28^. Here, we identify and characterize a ctenophore cytoplasmic IF protein—Cilin—that is phylogenetically distinct from canonical bilaterian cIFs and instead forms a well-supported clade with nematocilins and LMNTD proteins. These data clarify two important aspects of IF evolution: (1) the origin of cIFs predates the divergence of metazoan lineages, and (2) cilins represent a separate, lamin-derived IF lineage that followed a distinct evolutionary trajectory from the canonical cIFs that arose later in bilaterians. Our findings place the emergence of cIFs at the base of the animal tree and offer new insight into the structural innovations that accompanied the rise of multicellularity.

## Results

### Non-bilaterian animals typically encode two distinct intermediate filament genes

To understand the evolutionary history and origin of cIF genes, we conducted a wide phylogenetic analysis of their predicted protein sequences with emphasis on non-bilaterian animals. First, we identified IF proteins (lamins and cIF proteins) in publicly available genome assemblies, cDNA databases, and transcriptome assembly data using a sequence similarity search with BLAST followed by manual inspection. Since a central α-helical rod domain is the most conserved domain in all IF proteins, we used this domain for alignment and phylogenetic analysis^21^. Phylogenetic relationships of IFs proteins have been resolved by reconstructing phylogenetic trees using the Maximum likelihood approach. A Maximum Likelihood analysis was generated using translated amino acid sequences with the best-fit LG+R5 model (Supp. fig. 1). For this analysis a total of 120 IFs protein sequences were used representing all major animal groups (Supplementary file 1).

The majority of the analyzed non-bilaterian animal species possess two IF genes (Supp. Fig. 1; Fig. 2A). Our transcriptomic survey revealed only one IF gene in the crawling ctenophore *Coeloplana astericola* and the calcareous sponge *Leucetta chagosensis*. Interestingly, cnidarian *nematocilin* genes cluster together with some IF genes of Bilateria forming together a *nematocilin/cytovec* group (bootstrap support: 100) (Supp. Fig. 1; Fig. 2A). Like cnidarians, ctenophores also have IF genes forming two monophyletic clades. One branch of ctenophore IF genes clusters with the *nematocilin/cytovec* group (bootstrap support: 96) (Supp. Fig. 1), whereas the other ctenophore IF gene forms a separate group, likely corresponding to a *lamin* gene (Supp. Fig. 1; Fig. 2A).

**Figure 2.**
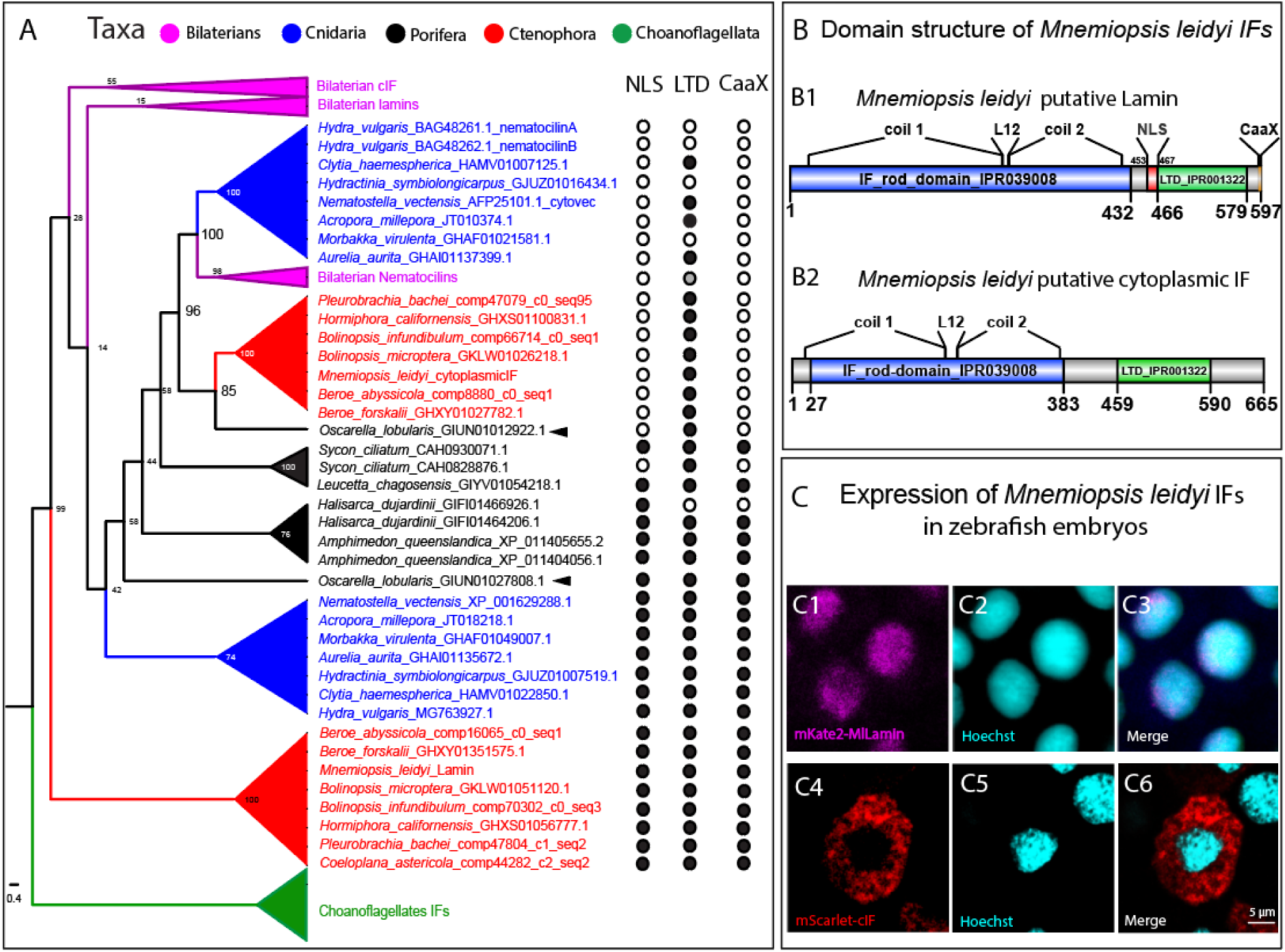
Evidence for cytoplasmic IF protein in ctenophores. **(A)** Collapsed cladogram of the IF phylogenetic tree from Supplementary Figure 1 and domain structure analysis of non-bilaterian IF proteins. Numbers at nodes are bootstrap values. Arrowheads indicate the phylogenetic positions of IF proteins from the sponge *Oscarella lobularis*. **(B)** Domain structure of *Mnemiopsis leidyi* IF proteins. **(C)** Expression of *Mnemiopsis leidyi* IF proteins in zebrafish embryos. **(C1–C3)** Intracellular localization of nuclear *Mnemiopsis leidyi* IF protein (lamin) in the cells of zebrafish embryo. **(C4–C6)** Intracellular localization of putative cytoplasmic *Mnemiopsis leidyi* IF protein in the cells of zebrafish embryo. CaaX – protein motif involved in targeting proteins to the endomembrane system; LTD – lamin tail domain; NLS – nuclear localization signal.

Most of the analyzed sponge species similarly possess two IF genes. However, in sponges, IF genes cluster by species, forming paraphyletic groups aligned with sponge classes and containing species-specific inparalogs. An exception is observed in *Oscarella lobularis* (Homoscleromorpha), whose IF genes appear separated on the phylogenetic tree (Supp. Fig. 1; Fig. 2A). Specifically, one IF gene sequence of *Oscarella lobularis* robustly clusters with ctenophore IF genes together the nematocilin/cytovec group (bootstrap support: 85), whereas the other sequence forms a distinct branch with the paraphyletic assemblage of poriferan IFs. Therefore, it is reasonable to suggest that the two IF genes identified in ctenophores do not belong exclusively to the lamin family. Instead, as observed in cnidarians, one gene probably encodes a nuclear lamin, while the other encodes a cIF protein.

### Domain architecture and intracellular localization support the presence of cytoplasmic IFs in ctenophores

To identify which IFs in non-bilaterian animals potentially localize in the cytoplasm, we analyzed their domain structure (Fig. 2A, B). Each sequence was examined for the presence or absence of a nuclear localization signal (NLS), a carboxy-terminal CaaX motif, and the lamin-tail domain (LTD). Our analyses revealed monophyletic group of IFs lacking the NLS and CaaX motif, suggesting they represent putative cIFs. This group includes cnidarian nematocilin sequences, non-canonical bilaterian cIFs, a clade of ctenophore nematocilin-like sequences, and a sequence from the sponge *Oscarella lobularis* (Fig. 2A). A NLS is, however, present in all other sponge sequences studied apart from *Halisarca dujardini* (Demospongiae). The presence or absence of the CaaX signal sequence mirrors the pattern observed for the NLS domain, with one exception: in *Amphimedon queenslandica* (Demospongiae), one sequence includes the NLS domain but lacks the CaaX motif.

No consistent pattern was observed for the LTD domain. This domain is missing in cnidarian nematocilins of *Hydra vulgaris*, *Hydractinia symbiolongicarpus*, and *Morbakka virulenta*, as well as in one sequence from the sponge *Amphimedon queenslandica*.

To investigate their subcellular localization, we tagged the two IF sequences from *Mnemiopsis leidyi* with fluorescent proteins and expressed them in zebrafish embryos (Fig. 2C). *M. leidyi* putative lamin localizes inside the nucleus. However, its distribution is dispersed rather than concentrated along the nuclear envelope (Fig. 2C1–С3). The putative cytoplasmic IF is found exclusively in the cytoplasm, where it is evenly distributed and forms short filaments, representing either single fibers or small aggregates (Fig. 2C4–С6).

Based on phylogenetic analyses of *Mnemiopsis leidyi* putative IF proteins, their domain prediction, and their heterologous localization in vertebrate cells, we conclude that ctenophores likely possess both intermediate filaments: nuclear IFs, such as lamin (Ml-Lamin), and cIF, which we designate as cilin (Ml-Cilin).

### Ctenophoran cIF protein, Ml-Cilin, is associated with cillia

To investigate the spatial distribution and subcellular localization of the identified IF proteins in *Mnemiopsis leidyi*, we developed custom antibodies targeting these proteins. One antibody was raised against the putative lamin (anti-Ml-Lamin, antigen E181–D409aa), whereas two were generated against the putative cIF protein, Ml-Cilin (anti-Ml-Cilin-C, antigen K22–E210aa; anti-Ml-Cilin-N, antigen M3–K115aa).

Immunohistochemistry experiments were conducted at the cydippid stage [2–3 days postfertilization (dpf)], a juvenile phase in *M. leidyi* life cycle (Fig. 3A). The Ml-Cilin protein was detected with high intensity signal in several regions. The subcellular localization depends on the structure: in comb plate cells Ml-Cilin is detected at the base of the cilia, in the aboral organ (ciliary grooves and balancers) within the entire cilia, and in cells at the base of the pharynx in the cytoplasm (Fig. 3B1–B3). In contrast, antibodies against the Ml-Lamin revealed a signal in the nuclei of all cells (Fig. 3C, D1–B3). The protein is distributed along the nuclear lamina, as it is typical distribution for lamins (Fig 3C). These findings demonstrate that ctenophores possess genuine lamins and cIF.

**Figure 3.**
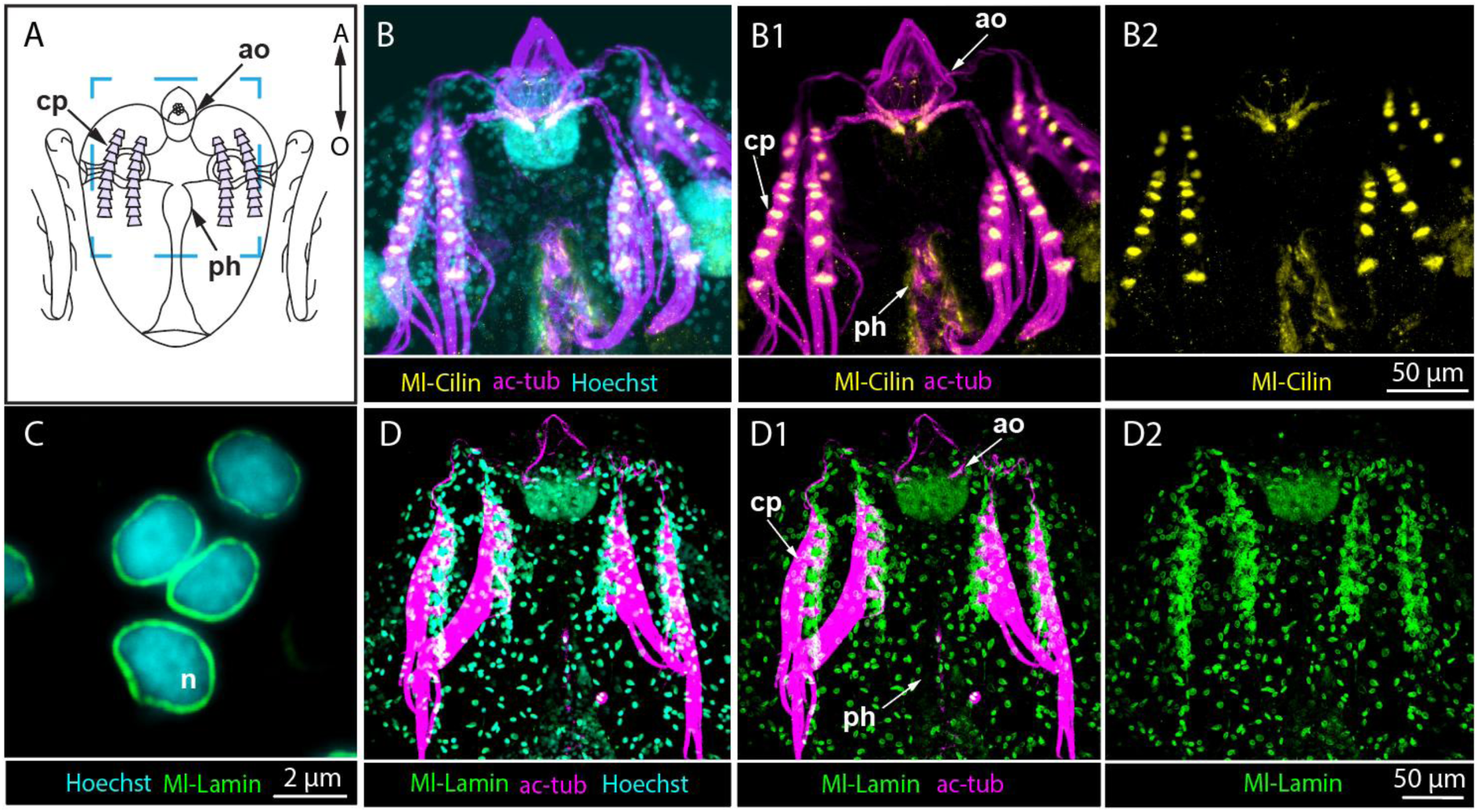
Localization of Ml-Lamin and Ml-Cilin, in the ctenophore *Mnemiopsis leidyi*. **(A)** Schematic drawing of a *Mnemiopsis leidyi* cydippid. Aboral is up. The blue square indicates the area shown in panels B–B2 and D–D2. **(B–B2)** Immunochemical staining reveals the localization of putative *Mnemiopsis leidyi* cytoplasmic IF, Ml-Cilin, in the aboral organ, comb plate, and pharynx. **(C–D2)** Immunochemical staining shows the localization of putative *M. leidyi* Lamin, Ml-Lamin, in the nucleus. Staining of cIF was performed using anti-Ml-Cilin-C antibodies. ac-tub – acetylated α-tubulin; A – aboral; ao – aboral organ; cp – comb plate; n – nucleus; O – oral; ph – pharynx.

Next, we conducted a more detailed analysis of Ml-Cilin localization. In the aboral organ (Fig. 4A), Ml-Cilin was observed at the base and along the length of the balancer cilia and the ciliary groove cilia (Fig. 4B1–B3). An even distribution of Ml-Cilin was detected in the cytoplasm of lithocytes. Additionally, in late cydippid stages (from 7 dpf) weak signal for Ml-Cilin can be seen at the base of dome cilia, which form a shielding structure over the aboral organ (Supp. Fig. 2C, C1). The second antibody, anti-Ml-Cilin-N, also detects Ml-Cilin in the cilia of the ciliary groove and balancer cilia in the aboral organ. However, in addition, the signal in the dome cilia is stronger than for the first antibody, anti-Ml-Cilin-C, and present at earlier stages, too (Supp. Fig. 2B, B1).

**Figure 4.**
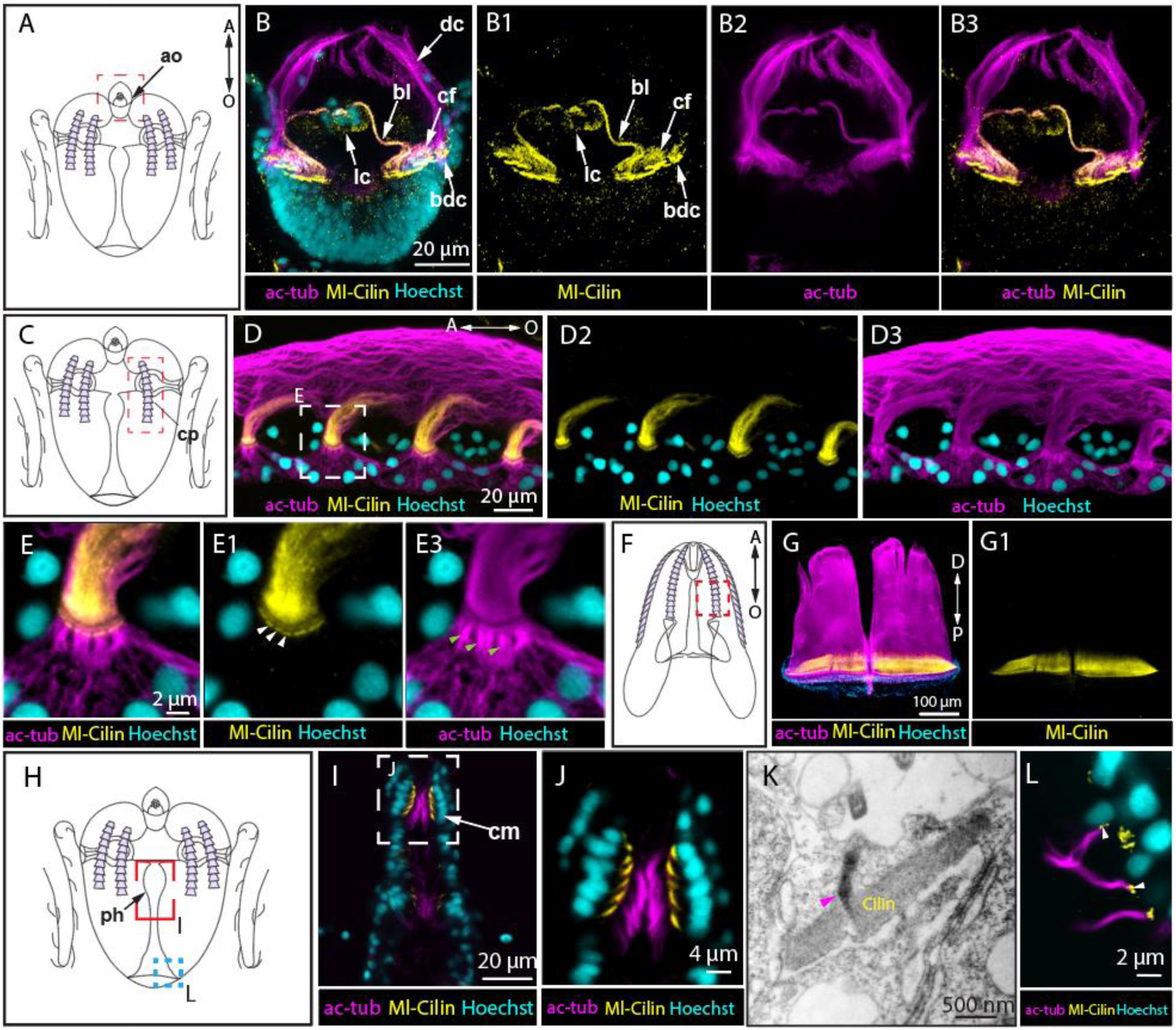
Ml-Cilin is associated with ciliary structures in *Mnemiopsis leidyi* cydippids and adults. **(A)** Schematic drawing of a cydippid showing the position of the aboral organ. **(B–B2)** Immunochemical double staining of the aboral organ for acetylated α-tubulin and Ml-Cilin. **(C)** Schematic drawing of a cydippid showing the location of the comb plates. **(D–E3)** Immunochemical double staining of comb plates for acetylated α-tubulin and Ml-Cilin. White arrowheads in E1 indicate intracellular Ml-Cilin structures localized at the base of acetylated α-tubulin bundles. Green arrowheads in E2 indicate intracellular bundles of acetylated α-tubulin. **(F)** Diagram of an adult (lobate stage) showing the location of comb plates. **(G, G1)** Immunochemical double staining of comb plates in lobate-stage for acetylated α-tubulin and Ml-Cilin. **(H)** Schematic drawing of a cydippid showing the pharynx area (I), mouth (L), gut (M). **(I, J)** Immunochemical double staining of the pharynx for acetylated α-tubulin and Ml-Cilin. **(K)** Transmission electron microscopy of Ml-Cilin bundles identified in the ciliary mill. Magenta arrowheads indicate ciliary rootlets. **(L)** Immunochemical double staining of cilia surrounding the mouth opening for acetylated α-tubulin and Ml-Cilin. White arrowheads indicate Ml-Cilin structures localized at the base of cilia, presumably centrioles. Staining of Ml-Cilin was performed using anti-Ml-Cilin-C antibodies. ac-tub – acetylated α-tubulin; A – aboral; ao – aboral organ; bdc – base of the dome cilia; bl – balancers; cf – ciliated furrow; cm – ciliary mill; cp – comb plates; dc – dome cilia; D – distal; lc – lithocytes; n – nucleus; O – oral; P – proximal; ph – pharynx.

In the cilia of the comb plates, Ml-Cilin was in the distal regions, with the strongest signal at the base of the cilia and a gradual decrease in signal intensity from distal to proximal direction (Fig. 4C, D1–D3). Beneath the apical surface of the cell, within the intracellular domain, we observed round structures positive for Ml-Cilin staining at the base of the ciliary rootlets, from which acetylated α-tubulin bundles extend (Fig. 4E– E3). These structures may represent microtubule-organizing centers (MTOCs) and are clearly detected in all replicates of the immunohistochemistry. However, they are not detected by anti-Ml-Cilin-N antibodies (Supp. Fig. 2D, D1).

In comb plates of adults (lobate stage, Fig. 4E), we also detected Ml-Cilin in the proximal region of the cilia at the base of the comb plates (Fig. 4F, G, G1, Supp. Fig. 2E, F, F1). The immunogold labeling visualized by transmission electron microscopy (TEM) enabled the precise localization of Ml-Cilin within the central region of the cilia in the comb plates (Supp. Fig. 3). The localization pattern of Ml-Cilin in the proximal region of the cilia within the comb plates closely mirrors the distribution of CTENO64, a structural protein responsible for linking adjacent cilia within the ciliary plate^29^. Double immunostaining against Ml-Cilin and CTENO64 demonstrated that their localization regions overlap almost completely (Supp. Fig. 4).

We also investigated in detail Ml-Cilin presence in pharyngeal cells of the ciliary mill (Fig. 4H). The ciliary mill consists of a dense arrangement of rigid cilia, which functions as a mechanical filter, breaking apart large particles and allowing only smaller ones to pass into the endodermal canal system^30^. Immunohistochemistry revealed a spindle-shaped structure beneath the cilia in the cytoplasm of the ciliary mill cells, angled inward into the cell (Fig. 4I, J). Using TEM, we were able to characterize this structure as a bundle of electron-dense filaments, embedding the rootlets of the cilia (Fig. 4K, Supp. Fig. 5A–F). The immunogold staining method demonstrated that these filamentous structures are positive for antibodies against *M. leidyi* Ml-Cilin (Supp. Fig. 5, G).

Structures of similar morphology were present in other ctenophores, including *Pleurobrachia* and *Beroe;* ^31,32^ in Beroe, these were confirmed as actin filament bundles through phalloidin staining. In view of these data, we performed a double staining for Ml-Cilin and F-actin, which showed overlapping signals (Supp. Fig. 6), suggesting that these bundle structures in *M. leidyi* pharynx cells are composed of both, Ml-Cilin and F-actin.

We also observed Ml-Cilin staining in cells with short cilia located around the mouth opening. In these cells, Ml-Cilin appear beneath the cilia in two rounded structures, whose shape and position suggest they are centrioles (Fig. 4L).

Since *M. leidyi* Ml-Cilin is strongly associated with ciliary structures across various ctenophore organs, we examined its localization in relation to sperm flagella. We therefore stained sperm cells of sexually mature ctenophores with anti-Ml-Cilin-C antibody. This revealed a distinct Ml-Cilin signal in the flagella of developing spermatids, as well as in a spherical structure adjacent to their nuclei (Fig. 5). High-magnification imaging showed that these spherical structures exhibit a toroidal morphology, with the spermatid flagellum anchored at their center (Fig. 5D). These findings suggest that Ml-Cilin is likely a structural component of the basal body of the sperm cell.

**Figure 5.**
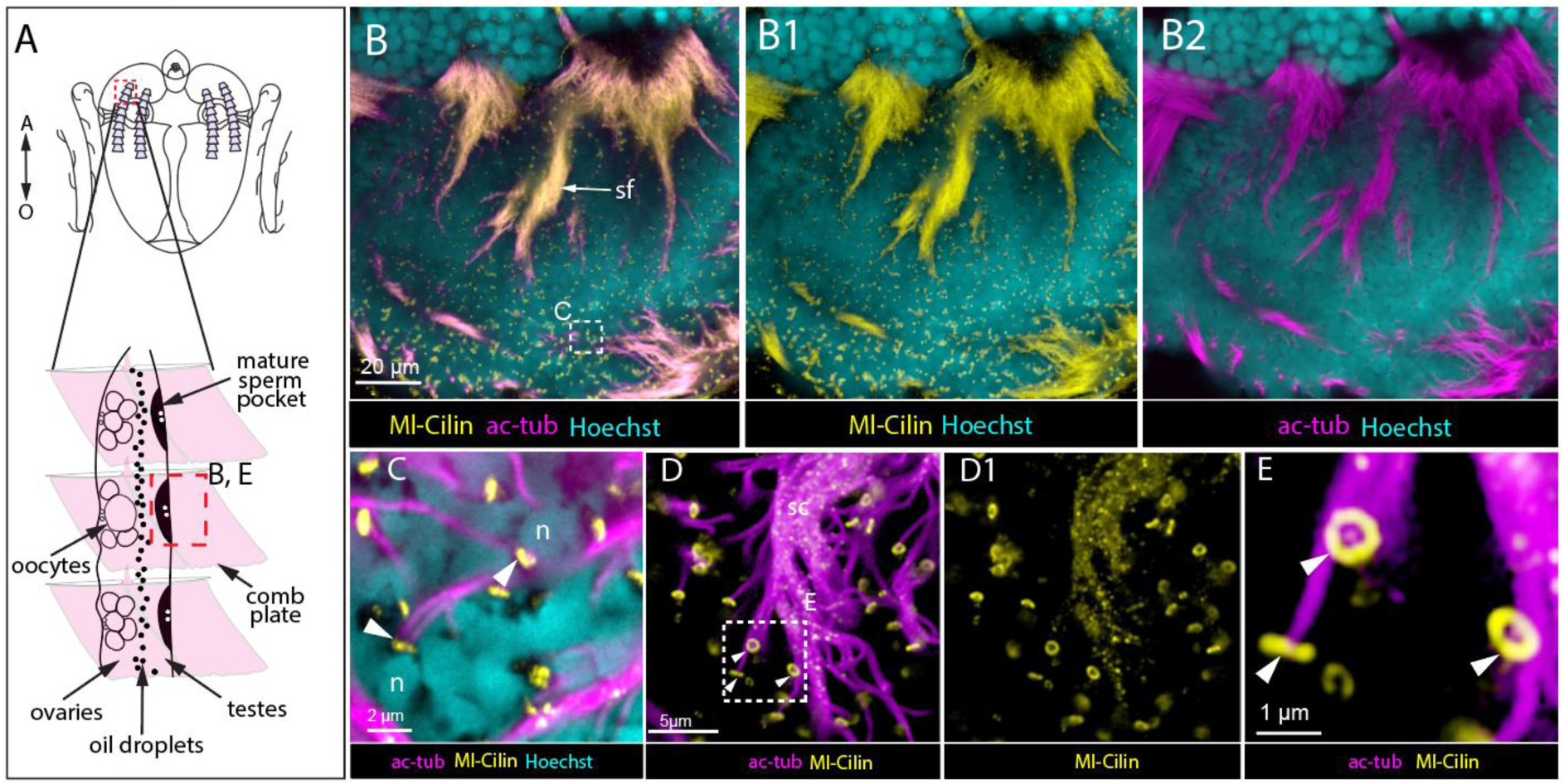
Ml-Cilin localizes in developing spermatids of *Mnemiopsis leidyi*. **(A)** Schematic drawing of a cydippid showing the position of the gonads. **(B–B2)** Immunochemical double staining of a mature sperm pocket containing developing spermatids using antibodies against acetylated α-tubulin and Ml-Cilin. **(C–E)** Immunochemical double staining of developing spermatids for acetylated α-tubulin and Ml-Cilin. White arrowheads indicate Ml-Cilin structures at the base of spermatid flagella. The Ml-Cilin signal at the base of the flagellum is most likely a structural component of the centriole. Ml-Cilin staining was performed using the anti-Ml-Cilin-C antibody. ac-tub – acetylated α-tubulin; A – aboral; n – nucleus; O – oral; sf – spermatid flagella.

Thus, Ml-Cilin in *M. leidyi* is consistently associated with ciliary structures across cydippid tissues, including the base and shaft of cilia in the aboral organ, comb plates, and pharyngeal ciliary mill, as well as in sperm flagella and basal body-like structures.

### Localization of ctenophoran cIFs beyond ciliary structures

In the following regions of 2 dpf cydippid we also identified Ml-Cilin not associated with cilia: tentacle bulb and the tentacle itself. A strong Ml-Cilin signal was detected in the outer cell layer of the tentacle bulb, just before its narrowing (Fig. 6B, B1, arrows). In these cells Ml-Cilin is uniformly distributed throughout the cytoplasm. Additionally, we observed Ml-Cilin in large cells within the tentacle bulb, which are likely myocyte progenitor cells^33^ (Fig. 6B–C). Furthermore, a distinct signal was detected in the inner cells of the tentacle, specifically in myocytes, organized into a continuous string (Fig. 6B, B1).

**Figure 6.**
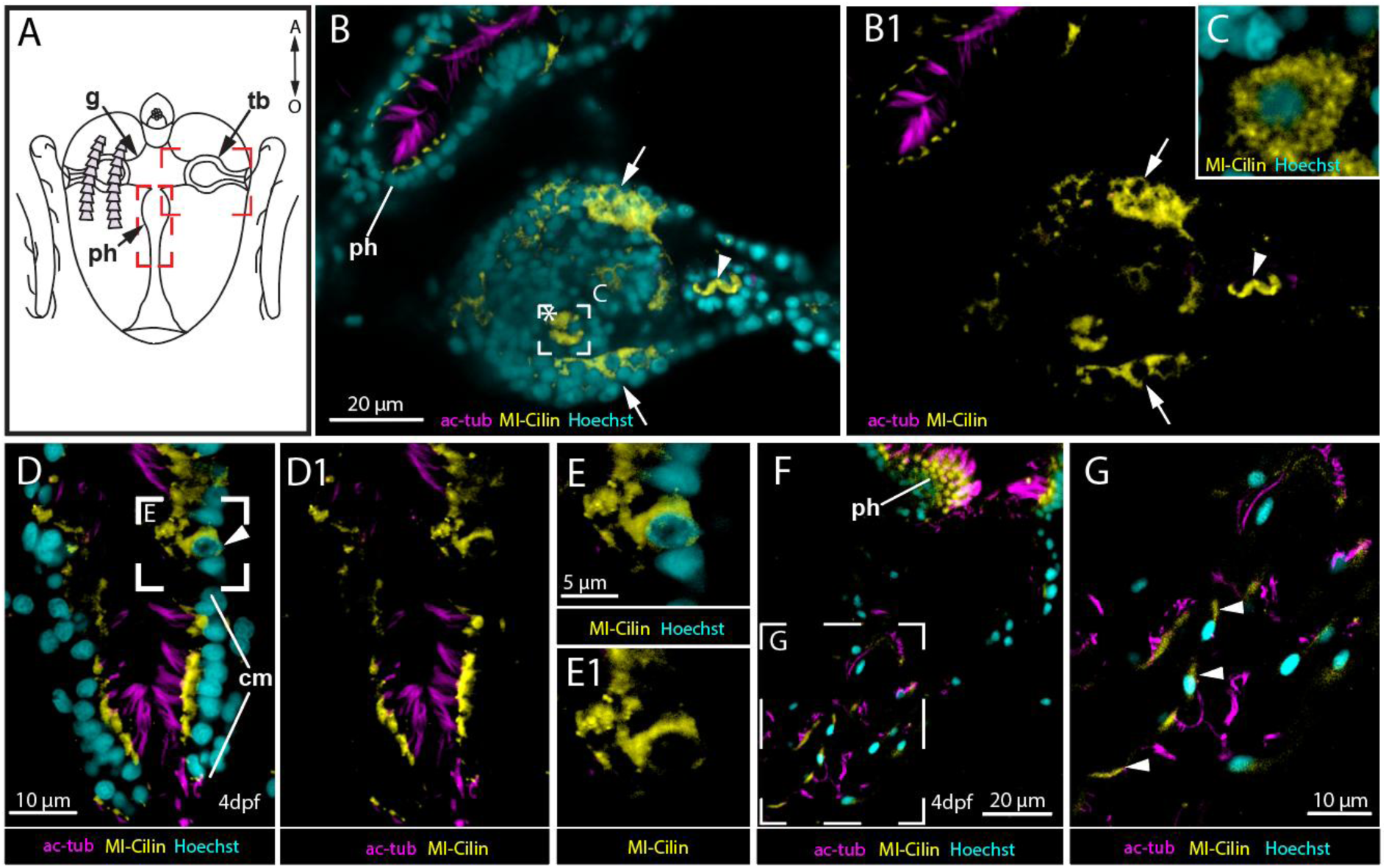
Ml-Cilin is also found outside ciliary structures in *Mnemiopsis leidyi*. **(A)** Schematic drawing of a cydippid showing the position of the pharynx, tentacle bulbs, and gut. **(B, B1)** Immunochemical double staining of the tentacle bulb for acetylated α-tubulin and Ml-Cilin. Asterisks mark muscle cell precursors. Arrows point to outer tentacle bulb cells positive for Ml-Cilin. Arrowhead points to muscle cells in the tentacle positive for Ml-Cilin. **(C)** Higher magnification of the same muscle precursor cell shown in B. **(D, D1)** Immunochemical double staining of the 4 dpf cydippid pharynx for acetylated α-tubulin and Ml-Cilin. Asterisks mark muscle cell precursors. Arrowhead points to a cell with Ml-Cilin evenly distributed throughout the cytoplasm. **(E, E1)** Higher magnification of the same pharynx cell shown in image D, with Ml-Cilin evenly distributed throughout the cytoplasm. **(F, G)** Immunochemical double staining of the 4 dpf cydippid gut for acetylated α-tubulin and Ml-Cilin. Arrowheads in G point to gut cells positive for Ml-Cilin. Staining of Ml-Cilin was performed using anti-Ml-Cilin-C antibodies. A – aboral; ac-tub – acetylated α-tubulin; g – gut; O – oral; ph – pharynx; tb – tentacle bulb.

As cydippid larvae develop, cIF signal appears in other tissues and cell types, still outside ciliary structures. By 4 dpf, we identified cells whose cytoplasm is uniformly filled with cIF. These cells are located in the pharynx, distal to the ciliary mill region (Fig. 6D–E1). At this location, an epithelial fold of the pharynx forms, separating the ciliary mill region from the rest of the pharynx. Additionally, we detected cIF staining in gut cells, where the cIF protein is distributed throughout the entire cytoplasm (Fig. 6F, G).

### Expression domains of cIF gene support immunostaining results

We performed *in situ* hybridization for the *Ml-cilin* gene using two different methods: hybridization chain reaction (HCR) and colorimetric whole mount *in situ* hybridization. These approaches were used to validate the Ml-Cilin protein localization data and to determine whether the protein is assembled and utilized in the same regions where its mRNA is processed. Both methods showed similar results, with intense expression in the comb plates, the ciliary mill region, the apical organ, and the tentacle bulb (Supp. Fig. 7A–F). Therefore, the patterns observed at the level of proteins are reproduced at the level of gene expression. This strengthens the reliability of the immunochemistry data and indicates a spatial correlation between gene expression and protein function.

### Bilaterian Ml-Cilin homologs are preferentially expressed in the ciliated cells of swimming larvae

Given the strong association of Ml-Cilin with ciliary structures, we examined whether bilaterian nematocilins, as Ml-Cilin homologs, show similar enrichment in ciliated cells. We analyzed single-cell RNA-seq datasets from trochophore larvae of the oyster *Magallana gigas* (Mollusca)^34^ and pluteus larvae of the sea urchin *Strongylocentrotus purpuratus* (Echinodermata)^35^, focusing on co-expression with conserved markers of ciliated cell identity (*tektin, rootletin,* and *foxj1*). In both species, larvae use specialized well-developed ciliary systems as an adaptation for effective locomotion and feeding in the planktonic environment. In *M. gigas* larvae, nematocilin expression was enriched in clusters of ciliated cells, with only background-level signal in non-ciliary cells (Figure 7A). In *S. purpuratus* larvae, nematocilin expression was likewise predominantly restricted to ciliated cell clusters, with only one non-ciliary cluster (cluster 6) showing expression slightly above the baseline mean value (Figure 7B). Taken together, these findings indicate that, similar to Ml-Cilin, the expression of bilaterian nematocilins is also associated with ciliated cells.

**Figure 7.**
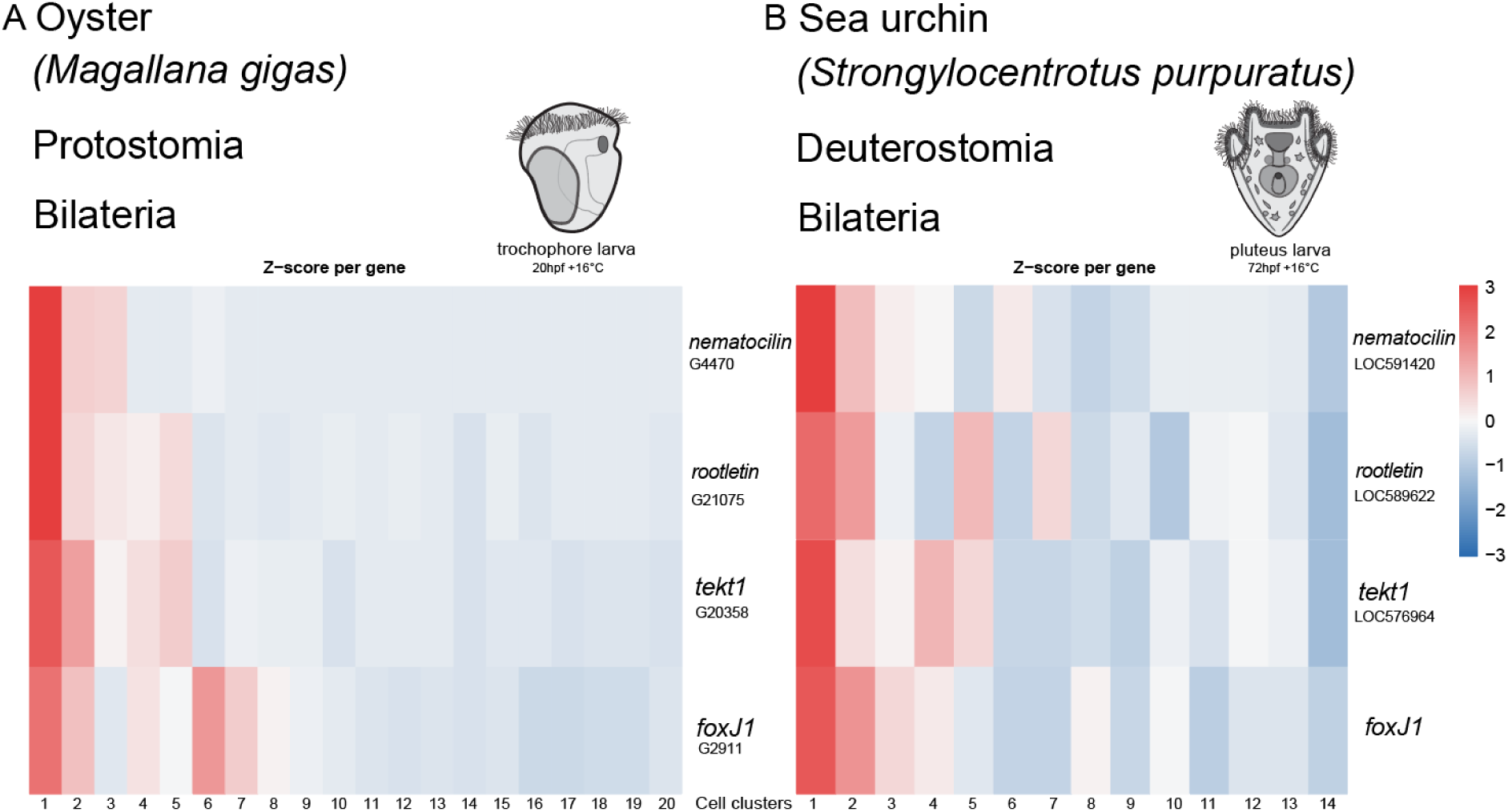
Expression of bilaterian nematocilins is enriched in clusters of ciliated cells in swimming larvae. **(A, B)** Heatmaps showing Z-score–normalized single-cell gene expression in **(A)** *M. gigas* trochophores and **(B)** *S. purpuratus* plutei.

### Vertebrate LMNTDs share homology and expression patterns with ctenophoran cIF, Cilin

In a 2015 study^7^, Kollmar reported that nematocilin is also present in humans, although in a truncated form lacking the α-helical rod domain. However, the study did not provide the full gene sequence or a database accession number. To investigate further, we extracted the human “nematocilin” sequence from the list of aligned sequences included in the publication and performed a BLAST search using NCBI tools. This search identified the gene *lamin-tail-domain-containing 1* (*LMNTD1*, gene ID: 160492). These proteins represent truncated forms of IF proteins, consisting of the C-terminal region that includes a lamin tail domain (LTD), while lacking the central α-helical rod domain, which is the most conserved region and typically involved in filament assembly.

In humans, two LMNTD genes, LMNTD1 (NCBI gene ID: 160492) and LMNTD2 (NCBI gene ID: 256329), exist (Supp. Fig. 8A) and encode multiple splice variants, as annotated in the NCBI database. Comparative analysis across vertebrates revealed that both genes are present in Mammalia, Lissamphibia, and Sauropsida. The exception within Sauropsida is that almost all birds retain only a single LMNTD gene (Supp. Fig. 8B–F). Among the mammalian species examined, only the echidna (*Tachyglossus aculeatus*) retained an N-terminal α-helical segment in the LMNTD gene, though markedly shortened (Supp. Fig. 8B). In lizards, a similar pattern is observed. LMNTD2 is rodless, and LMNTD1 contains an α-helical segment that is significantly shorter, roughly half the length of that found in turtles and frogs (Supp. Fig. 8E). In turtles and frogs, LMNTD2 lacks the rod domain, whereas LMNTD1 maintains the full IF architecture (Supp. Fig. 8D, F). Interestingly, although the α-helical segment in LMNTD1 resembles the canonical IF rod domain, it is not recognized by either InterPro or the NCBI Conserved Domain Database (CDD), suggesting a degree of divergence. In the ray-finned fish *Acipenser ruthenus* (sterlet), an LMNTD-like protein with IF-like structural features is also present (Supp. Fig. 8G), although its rod domain remains also undetected by InterPro and CDD tools. Notably, no LMNTD homologs were identified in teleost fishes, suggesting lineage-specific loss of these genes. In contrast, LMNTD-like proteins in both lungfish and a cartilaginous fish species retain clearly identifiable IF rod domains and LTD regions, supporting their structural conservation in these lineages (Supp. Fig. 8H, I). The three-dimensional structures of these proteins were modeled using AlphaFold2 (Supp. Fig. 8).

To explore whether LMNTD genes are truly related to nematocilins and ctenophore cilins, we built a phylogenetic tree using the N-terminal regions of the proteins, specifically those that include an α-helical segment. The bird and human LMNTD gene sequences were excluded from the analysis due to the short and unstructured N-terminal ends of their LMNTD proteins (Supp. Fig. 8A, С). Our analysis showed that all LMNTD proteins form a monophyletic group, robustly nested within the nematocilin/cytovec/cilin clade, and are sister to the nematocilin of ascidian *Ciona intestinalis* (bootstrap support value: 96; Supp. Fig. 9).

Due to the loss of the highly conserved N-terminal α-helical segment in LMNTD proteins of birds and most mammals, including humans, these sequences cannot be reliably included in phylogenetic analyses. To assess whether these truncated proteins are truly homologous to LMNTD family members, we conducted BLASTP searches using LMNTD sequences from the nearest available species against the non-redundant (nr) protein database. A BLASTP search using the turtle (*Emys orbicularis*) LMNTD1 sequence (XP_065279841.1) against the duck (*Anas platyrhynchos*) proteome identified LMNTD1 (XP_021130469.1) as the top hit (E-value: 7e-127), followed by lamin (XP_005021763.4) with a significantly weaker match (E-value: 7e-16). Similarly, querying the mammal, echidna (Tachyglossus aculeatus), LMNTD1 sequence (XP_038597040.1) against the human proteome returned LMNTD1 (Homo sapiens, XP_054227227.1) as the highest-scoring hit (E-value: 8e-65), followed by lamin B3 (Homo sapiens, AIZ72720.1) with an E-value of 9e-11. These results, combined with phylogenetic analysis, support the homology of LMNTD proteins with nematocilin and ctenophore cilins and suggest a shared evolutionary origin.

To explore the cell-type specificity of human LMNTD gene expression, we analyzed single-cell RNA sequencing data available in the Human Protein Atlas database^36^. This resource integrates publicly available genome-wide expression data from 40 human tissues, encompassing 81 distinct cell types. Using this dataset, we found that LMNTD1 is enriched in early spermatids, late spermatids, ciliated cells, and astrocytes. Among ciliated cells, LMNTD1 is expressed in the endometrium, fallopian tubes, lungs, and bronchi (Supp. Fig. 10A). In contrast, LMNTD2 is enriched exclusively in early and late spermatids (Supp. Fig. 10B). Thus, similar to ctenophore Ml-Cilin, human LMNTD proteins are primarily associated with ciliated cells and sperm cells.

## Discussion

In this study, we present compelling evidence for the presence of a cytoplasmic intermediate filament (cIF) in *Mnemiopsis leidyi*, a comb jelly from the clade Ctenophora, the sister group to all other metazoans^28^. We designate this filament as Ml-cilin. Based on a combination of phylogenetic analyses, domain structure, and subcellular localization, our findings suggest that cIFs emerged in the lineage to the last common ancestor of metazoans. Supporting this, we found no orthologs of ctenophoran cIFs in choanoflagellates, the closest unicellular relatives of animals^37,38^, indicating that cIFs are a metazoan-specific innovation. Likely derived from an ancestral lamin through gene duplication and functional divergence, *M. leidyi* Ml-Cilin, together with its homologs across diverse metazoan lineages, defines a distinct class within the IF protein family.

Our phylogenetic analysis supports the hypothesis that ctenophore cilins are orthologous to nematocilins previously identified in cnidarians and bilaterians^7^, suggesting a shared evolutionary origin. Notably, we found that the vertebrate *LMNTD* genes correspond to these previously described nematocilin orthologs. These genes not only share sequence similarity with nematocilins and ctenophore cilins but also display remarkably similar expression patterns. Specifically, LMNTD1 and LMNTD2 are predominantly expressed in ciliated cells and spermatids, closely mirroring the localization patterns of Ml-Cilin in *M. leidyi*. Based on these findings, we propose grouping ctenophore cilins, cytovec of *Nematostella*, nematocilins of other cnidarians and bilaterians, and LMNTD proteins under the unifying name Cilin.

The spatial distribution of Ml-Cilin in *M. leidyi* reveals its strong association with ciliary structures, including the comb plates, aboral organ, ciliary grooves, and the pharyngeal ciliary mill. In these regions, Ml-Cilin consistently co-localizes with acetylated α-tubulin and structural components such as CTENO64^29^, suggesting a role in stabilizing ciliary rootlets and supporting coordinated ciliary motion. Its localization in the basal bodies of spermatids further points to a crucial role in flagellar organization.

At the same time, Ml-Cilin is also expressed in non-ciliated tissues, including myocyte precursors in the tentacle bulb and differentiated muscle cells, where it shows uniform cytoplasmic distribution. Notably, in vertebrates, maintaining the integrity of muscle cells and their function requires structural support from canonical cIFs, particularly desmin, which helps maintain the alignment of myofibrils^39^. This similarity raises the possibility that, despite distinct evolutionary origins of desmin and ctenophore cIF, ctenophore cIF may serve a comparable structural role, convergently contributing to muscle cell organization in ctenophores.

Taken together, these findings suggest that Ml-Cilin in *M. leidyi* may serve dual roles, both in ciliary architecture and broader cytoskeletal functions. This multifunctionality could be achieved through several mechanisms, including alternative splicing and post-translational modifications, both of which are well-established strategies for regulating cIFs in bilaterian animals^40–42^. Since Ml-Cilin is observed in multiple tissues and cell types in *M. leidyi*, it is reasonable to assume that regulatory mechanisms allowing a single cIF to perform diverse functions were already present in early metazoans. However, these possibilities remain to be explored in detail by future studies.

Considering all the data, we propose a revised scenario for the evolution of intermediate filaments. In this view, cilin—the first cIF—arose in the common ancestor of animals through a duplication of lamin, followed by neofunctionalization of one of the copies (Fig. 8, event 1). Subsequently, in the bilaterian ancestor, a second independent duplication of lamin occurred, and neofunctionalization of one of the resulting copies led to the emergence of canonical cIFs (Fig. 8, Event 2). Later, cilin duplicated again in chordates, and one of the paralogs lost the rod domain, giving rise to a truncated variant of cilin (Fig. 8, Event 3). The coexistence of both full-length cilin and its truncated paralog can be observed in *Xenopus tropicalis* (Supp. Fig. 8F). In some vertebrate lineages, such as mammals, both cilin paralogs lost the rod domain, resulting in two truncated cilin proteins—LMNTD1 and LMNTD2—as seen in Homo sapiens (Fig. 8, Event 3). We propose that the evolution of IF involved at least two major episodes of lamin duplication and neofunctionalization, giving rise to the cilin lineage—comprising cilin proteins, and the bilaterian-specific canonical cIF proteins (Fig. 8).

**Figure 8.**
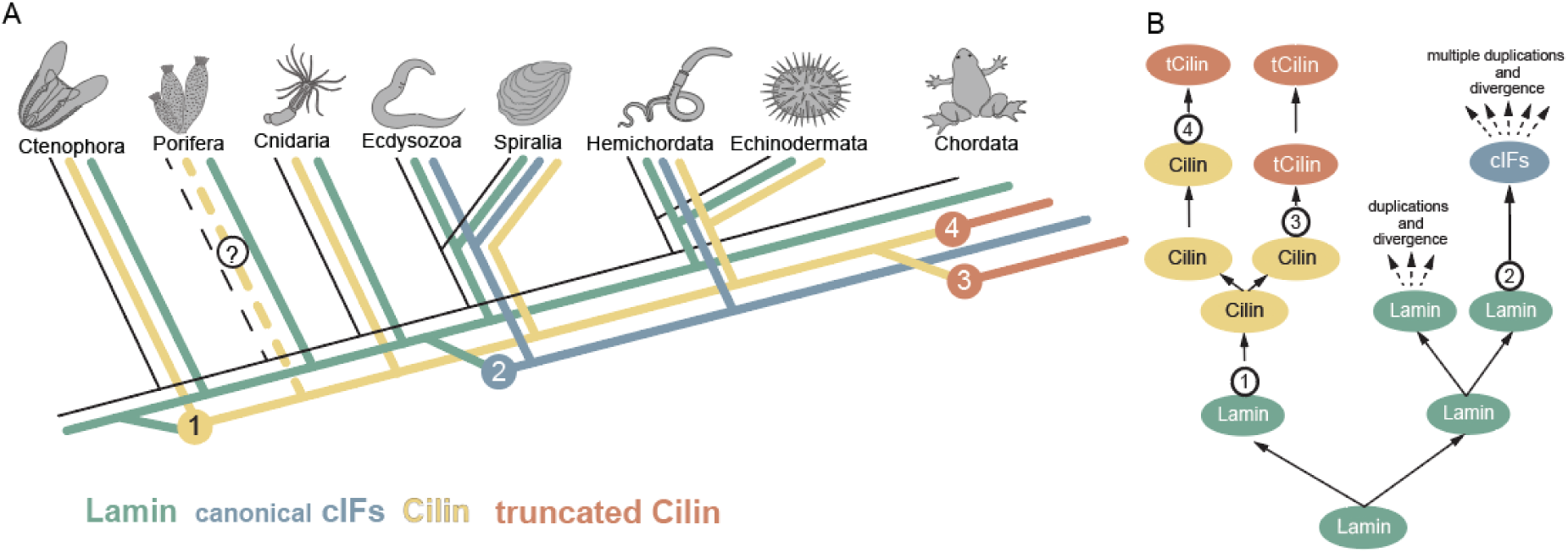
Proposed evolutionary scenario for IFs in Metazoa. **(A)** Schematic phylogenetic tree of metazoan lineages illustrating key events of lamin and IF gene duplication followed by neofunctionalization. **(B)** Stepwise model of IF evolution: Event 1: a lamin duplication in the common ancestor of ctenophores and other metazoans gave rise to cilin, the first cIF. Event 2: An independent lamin duplication and neofunctionalization in the common ancestor of bilaterians led to the emergence of canonical cIFs. Event 3: in the vertebrate lineage, cilin duplicated, and one copy lost the rod domain, giving rise to the truncated form of the cilin. Event 4: In mammals, a second cilin copy also lost the rod domain, resulting in two paralogous truncated cilins.

The localization of Cilin to cilia and sperm flagella in *Mnemiopsis leidyi*, along with the conserved expression of its orthologs Cilin1 (aka LMNTD1) and Cilin2 (aka LMNTD2) in human ciliated cells, suggests an evolutionary ancient role in ciliary structure and function (Fig. 9). Additionally, the presence of Cilin1 at the centrosomes of human cell lines RT-4 and SH-SY5Y, as documented in the Human Protein Atlas database^43^, supports its potential role in MTOC or ciliary organization. These data together imply that cilia-associated IFs have been functionally conserved early in animal evolution. This evolutionary continuity implies that ciliary dysfunctions seen in clinical practice, particularly those causing infertility or ciliary epithelium disorders, may reflect disruptions in ancient cellular architecture inherited from the first multicellular animals. Despite this apparent ancestral role, cilin appears to have been lost in some lineages for example Ecdysozoa^7^ or Teleostei, possibly due to functional redundancy with other cIF proteins or lineage-specific shifts in ciliary architecture and its regulation.

**Figure 9.**
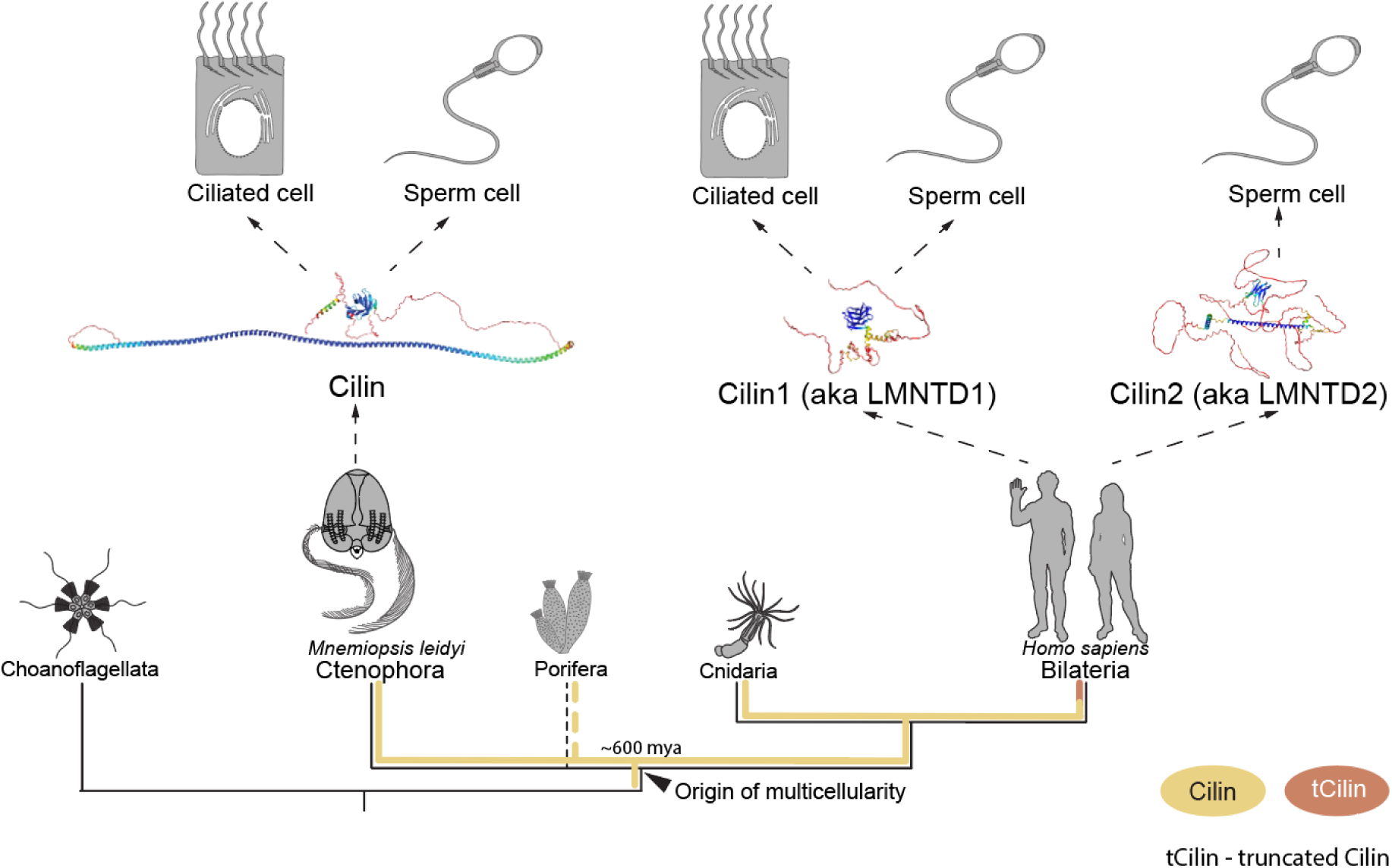
Proposed evolutionary and functional conservation of ctenophore and human cilins. Schematic representation of the conserved localization and functional roles of *Mnemiopsis leidyi* and human cilin in ciliated and sperm cells. The shared expression patterns support functional conservation of cilia-associated intermediate filaments inherited from the last common ancestor of metazoans.

The IF protein cilin in *Mnemiopsis leidyi* exhibits extensive spatial and functional interactions with other cytoskeletal networks, including actin filaments and microtubules. In particular, its co-localization with F-actin bundles in the pharyngeal ciliary mill and with acetylated α-tubulin in ciliary rootlets and axonemes points to a key structural role in coordinating cytoskeletal dynamics. Such integration is critical for maintaining cellular integrity by modulating the mechanical properties of composite cytoskeletal structures, thereby enhancing the ability of cells to resist mechanical stress^44^. These observations underscore the importance of identifying the molecular crosslinkers or adaptor proteins that mediate interactions between cytoskeletal systems. Proteins such as plectins^45^ or APC^46^ are known to mediate interactions of сIFs, actin, and tubulin in vertebrates^47^. However, nothing is currently known about how and when such cross-cytoskeletal interaction systems first emerged in evolution. Based on our findings, we propose that this integrative cytoskeletal network may have emerged early in metazoan evolution, or possibly even prior to the origin of multicellularity. Deciphering these components in *M. leidyi* could reveal conserved strategies for assembling complex, interconnected cytoskeletal architectures, vital for the evolution of multicellularity and tissue specialization.

Exploring how cilin integrates into ancestral cellular systems offers a rare window into the structural innovations that preceded the diversification of animal forms. As genomic and proteomic data accumulate across various metazoan lineages, from ctenophores to vertebrates, comparative analyses may reveal whether cilin contributed to fundamental features of metazoan evolution, such as increasing tissue complexity, the capacity to integrate and transmit mechanical forces across multicellular architectures or stabilization of ciliary architecture and function. Such findings could fundamentally revise our understanding of how cytoskeletal systems co-evolved with the rise of multicellularity in animals.

## Materials and Methods

### Mnemiopsis leidyi culture

Full life cycle of *M. leidyi* based on cydippid reproduction was established in the lab. Groups of 20–40 individuals were kept in 3-liter Kreisel tanks at 17–19 °C, with a salinity of 27.5 ppt artificial seawater (ASW) and a 17:7 light/dark cycle. The animals were fed once daily with the rotifer *Brachionus plicatilis*, which had been cultured on a commercial concentrated microalgae solution, RG Complete (Reef Nutrition – Reed Mariculture, California, USA). For embryo collection, 10–20 individuals were transferred to 200–250 mL beakers, where they spawned daily under these conditions.

### Phylogenetic analyses

The *Nematostella vectensis* cIF gene *cytovec* (accession number AFP25101.1) was used as a query for a tblastn search against the *Mnemiopsis leidyi* genome via the *Mnemiopsis* Genome Project Portal (https://research.nhgri.nih.gov/mnemiopsis/). Candidate sequences identified were analyzed for the presence of the conserved intermediate filament rod domain and the lamin tail domain using InterPro (https://www.ebi.ac.uk/interpro/). The retrieved *M. leidyi* sequences were then used as queries in additional tblastn searches on the NCBI TSA database (excluding *M. leidyi*) and the NeuroBase database (https://neurobase.rc.ufl.edu/pleurobrachia/blast?view=blast) to identify putative intermediate filament sequences from other ctenophore species. In total, 120 amino acid sequences from choanoflagellates, ctenophores, cnidarians, and bilaterians were included in the final analysis.

Phylogenetic analysis was carried out separately for two IF domains: the intermediate filament rod domain and the lamin tail domain. These domains, InterPro IPR039008 (intermediate filament rod) and InterPro IPR001322 (lamin tail domain), were identified using the InterPro database (https://www.ebi.ac.uk/interpro/). Amino acid sequences were aligned using the MUSCLE algorithm (v3.8.31)^48^. Poorly aligned regions were filtered out using TrimAL (v1.2rev59)^49^ with the “automated1” heuristic setting, which selects the optimal trimming strategy automatically. The alignments were used without manual modification. Phylogenetic reconstruction was conducted using the maximum likelihood method implemented in IQ-TREE (v2.0-rc2)^50^, applying the LG + R6 substitution model, which was identified as the best fit for the dataset by ModelFinder Plus (MFP). Node support was evaluated using 1000 bootstrap replicates. The resulting phylogenetic tree was visualized with FigTree (v1.4.4)^51^, and final graphical adjustments were made in Adobe Illustrator (version 29.6; Adobe Inc., San Jose, CA, USA), no changes were made to the tree topology or branch lengths.

### Signaling motifs analysis

Each sequence was screened for the presence of a nuclear localization signal (NLS motif). NLS motifs were predicted using the NucPred web server^52^, cNLS Mapper^53^ (cut-off score = 2.0) and DeepLoc - 2.1^54^. To identify the presence of a CaaX motif, the PrePS tool^55^ was used.

### Single-cell RNA sequencing data analysis

Single-cell RNA sequencing datasets were obtained from the NCBI Gene Expression Omnibus (GEO): *Magallana gigas*^34^ trochophore larvae (GSM7296642)^35^ and sea urchin (*Strongylocentrotus purpuratus*)^35^ pluteus larvae at 72 hpf (GSM4704281). Filtered feature-barcode matrices (matrix.mtx, barcodes.tsv, features.tsv) provided with each dataset were used as input. Data preprocessing and analysis were performed in R (v4.5.1) using the Seurat package (v4.0). Gene identifiers were taken directly from the feature tables of each dataset.

Average gene expression across clusters was calculated with Seurat’s AverageExpression function. For visualization, values were z-score normalized across clusters to highlight relative differences in expression. Heatmaps were generated using the pheatmap package, with candidate marker genes of interest (e.g., LOC591420, LOC576964, LOC589622, FoxJ1 for S. purpuratus; G4470, G2911, G20358, G15286 for M. gigas). Clusters were optionally reordered to emphasize biological comparisons.

### mRNA microinjections

For overexpression experiments of *M. leidyi* intermediate filament proteins in zebrafish embryos, the full-length coding sequences of *Ml-cilin* (gene model ML08883a) and *Ml-lamin* (gene model ML32592a) were cloned into the pCRII-mScarlet3 and pCRII-mKate2 vectors (pCRII vector, Thermo Scientific, Waltham, MA, USA), respectively, each flanked by an SP6 promoter and an SV40 polyadenylation signal. This resulted in N-terminally fluorescently tagged recombinant proteins. The resulting constructs were linearized using EcoRV, and capped mRNA was synthesized using the SP6 mMESSAGE mMACHINE Kit (Cat# AM1340, Invitrogen, Thermo Fisher Scientific, Waltham, MA, USA). For each embryo, 600 pg of mRNA was injected at the one-cell stage. Phenol red (0.05 %) (Cat# P0290, Sigma-Aldrich, Burlington, MA, USA) was included in the injection mix as a tracer.

### Immunohistochemistry

Custom rabbit polyclonal antibodies against *M. leidyi* anti-Ml-Lamin and anti-Ml-Cilin-C proteins were produced by ABclonal (ABclonal, Inc., Woburn, MA, USA). The anti-Ml-Lamin antibody was generated using a recombinant protein corresponding to amino acids E181–D409 of the Ml-lamin sequence, expressed in E. coli Rosetta cells using pET-28a and pET-28a-SUMO vectors with N-terminal His, T7, and SUMO tags. The antigen was purified and used to immunize Japanese White rabbit. Following four rounds of immunization, antigen-specific affinity purification was performed, yielding antibody concentrations of 1.24 mg/mL. The anti-Ml-Cilin-C antibody was similarly generated using a recombinant protein corresponding to amino acids K22–E210 of the cIF protein. The antigen was also expressed in E. coli Rosetta using the same vector system and tags, then purified for immunization of New Zealand White rabbit. Affinity purification produced antibodies at concentrations of 1.28 mg/mL. Both antibodies were supplied in PBS (pH 7.3) containing 50% glycerol and 0.05% ProClin 300 and demonstrated positive antigen recognition via ELISA and Western blot.

Additional rabbit anti-Ml-Cilin-N antibodies were generated against GST-tagged truncated recombinant Ml-cilin (antigen: N-terminal, M3–K115 aa; #IG-1452/2220-5.6; conc. 81 μg/ml; immunoGlobe GmbH, Himmelstadt, Germany). After immunization and six additional boosts, the blood serum was affinity purified on tandem arrays of GST and truncated Ml-cilin columns.

For immunostaining, *M. leidyi* cydippids were fixed as follows: samples were prefixed with Rain-X (ITW Global Brands division of Illinois Tool Works Inc., Houston, TX, USA) (200 µL per 1 mL of sample) for 15 minutes, then washed three times for 10 minutes each in PBTx (0.2 % Triton X-100 in PBS; 137 mM NaCl, 2.68 mM KCl, 10.14 mM Na₂HPO₄, 1.76 mM KH₂PO₄, pH 7.4)^56^. The samples were then rapidly transferred to ice-cold 100 % methanol and stored at –20 °C for 1 to 3 hours (not exceeding 3 hours). After methanol fixation, samples were rehydrated by washing five times for 5 minutes in PBTx. Rehydrated cydippids were incubated for 1–2 hours at room temperature on a shaker in blocking solution (5–10 % sheep serum and 1 % BSA (Cat# A-7906. Sigma-Aldrich, Burlington, MA, USA) in PBTx). Custom antibodies were used in combination with mouse acetylated α-tubulin antibody (T6793, Sigma-Aldrich, Burlington, MA, USA) to label ciliary structures. All primary antibodies were diluted in blocking solution, centrifuged at 16,000 × g for 10 minutes, and the supernatant was used for overnight incubation at 4 °C. Final concentrations were 2–4 µg/mL for custom *M. leidyi* antibodies and 1 µg/mL for acetylated α-tubulin. For samples undergoing double staining with antibody and conjugated phalloidin–Alexa Fluor Plus 555 (Cat# A30106; Invitrogen, Thermo Fisher Scientific, Waltham, MA, USA), cydippids were fixed for 30 minutes on ice in 4% PFA (Cat# 1.04005.1000; Merck KGaA, Darmstadt, Germany) in PBS, then immediately processed for immunostaining. Phalloidin was used at a 1:200 dilution from a 1 unit/μL stock solution (equivalent to 1000 units/mL or ∼6,600 nM), resulting in a final concentration of 5 units/mL (∼33 nM). It was added to the secondary antibody solution to allow simultaneous visualization of F-actin structures.

Following incubation with primary antibodies, samples were washed five times for 10 minutes each in PBTx. Secondary antibodies (anti-rabbit and anti-mouse, Alexa Fluor Plus 488 and 594 conjugated, Cat# A32731, Cat# A48288, Invitrogen, Thermo Fisher Scientific, Waltham, MA, USA) were diluted 1:250 in blocking solution, centrifuged at 16,873 × g for 10 minutes, and the supernatant was used for staining. Hoechst 33342 (TA9H97BAECD2, Targetmol Chemicals Inc. USA) was added to the secondary antibody mix at a final concentration of 1 µg/mL for DNA staining. Samples were incubated overnight at 4 °C with secondary antibodies, then washed three times for 15 minutes in PBTx, followed by five washes in PBS. Finally, samples were mounted on glass slides using Vectashield (Vector Laboratories, Inc., USA) or 80 % glycerol and imaged with a Carl Zeiss LSM 980 (Carl Zeiss Microscopy GmbH, Jena, Germany). To enhance visualization of the regions of interest, surrounding areas were excluded using clipping plane tools, and image contrast was optimized by adjusting brightness, gamma, and intensity thresholds. All adjustments were applied to the entire image, not locally. Image processing was performed using ImageJ (Version 1.54g)^57^ and Adobe Photoshop (Version 26.8.1; Adobe Inc., San Jose, CA, USA).

To verify antibody specificity, we performed a preadsorption control by incubating the diluted primary antibodies with 1 µg/mL of the corresponding recombinant antigen used for immunization (expressed in E. coli and purified via affinity chromatography). This step is expected to block specific staining. Preincubation of anti-Ml-Cilin-C (antigen: E181–D409 aa) and anti-Ml-Lamin (antigen: K22–E210 aa) antibodies with their respective antigens resulted in complete loss of signal, confirming staining specificity (Supplementary Figure 11, A–D1).

### Western blot

*Mnemiopsis leidyi* tissues were incubated in 100 μl of Cell Extraction Buffer (ThermoFisher, FNN0011) supplemented with protease inhibitors (Roche Diagnostics GmbH, Mannheim, Germany) to prepare lysates. Samples were centrifuged at 16,000 × g for 15 minutes at 4 °C, and the resulting supernatants were mixed with 10 μl of 6хLaemmli SDS sample buffer (Cat# J61337-AD, Thermo Scientific, Waltham, MA, USA). Proteins were separated by SDS-PAGE and transferred to nitrocellulose membranes. Membranes were blocked using 10 % skimmed milk dissolved in PTw (1× PBS, pH 7.4, containing 0.1 % Tween-20 (Cat# P9416, Sigma-Aldrich, Burlington, MA, USA)). Primary antibodies against anti-Ml-Lamin, anti-Ml-Cilin-C and β-actin (#4970S, Cell Signaling Technology, Inc., Danvers, MA, USA), were diluted 1:1000 in blocking solution and incubated with the membranes overnight at 4 °C. After washing with PTw, membranes were treated with an HRP-conjugated anti-rabbit IgG secondary antibody (1:100,000; Cat# A0545, Sigma-Aldrich, Burlington, MA, USA). Signals were visualized using the SuperSignal West Femto Maximum Sensitivity Substrate (Cat# 34094, Thermo Scientific, Waltham, MA, USA).

### In situ hybridisation

The colorimetric *in situ* hybridization procedure for *Mnemiopsis leidyi* was adapted from a previously established protocol developed for the sea anemone *Nematostella vectensis*^58^. Specimens were initially fixed on ice in a solution of 3.7 % formaldehyde and 0.2 % glutaraldehyde in PBS for 15 minutes. This was followed by an additional fixation step in 3.7 % formaldehyde in PBS for one hour at room temperature with gentle agitation. After fixation, samples were washed three times for 15 minutes each in PTw buffer (1× PBS containing 0.1 % Tween-20, pH 7.4), then gradually dehydrated through increasing concentrations of methanol before being stored in 100 % methanol at –20 °C. Digoxigenin (DIG)-labeled RNA probes were generated from clones containing the full-length *Ml-cilin* coding sequence (gene model ML08883a), subcloned into the pJet1.2 vector (Cat# K1232, Thermo Fisher Scientific, Waltham, MA, USA), using the MEGAscript™ SP6 Transcription Kit (Cat# AM1330, Invitrogen, Thermo Fisher Scientific, Waltham, MA, USA). Prior to hybridization, fixed cydippids were rehydrated through a descending methanol series (60 % and 30 % methanol in PTw), treated with 80 µg/mL Proteinase K in PTw for 40 seconds, and then washed in 2 mg/mL glycine in PTw. Hybridization was carried out overnight at 58 °C using DIG-labeled RNA probes diluted to 0.5 ng/mL in hybridization buffer (50% formamide, 5× SSC [0.75 M NaCl, 0.075 M sodium citrate, pH 4.5], 50 µg/mL heparin, 500 µg/mL torula yeast RNA, 1% SDS, 0.1% Tween-20, nuclease-free water). Detection of bound probes was achieved using anti-DIG-AP Fab fragments (Cat# 11093274910, Roche Diagnostics GmbH, Mannheim, Germany) at a 1:2000 dilution in 0.5 % blocking reagent (Cat# 11096176001, Roche Diagnostics GmbH, Mannheim, Germany) in 1× MAB buffer (100 mM maleic acid, 150 mM NaCl, pH 7.5). Signal development was performed using a chromogenic reaction with NBT(Cat# 11383213001)/BCIP(Cat# 11383221001) substrate (Roche Diagnostics GmbH, Mannheim, Germany).

Hybridization Chain Reaction (HCR) was carried out based on the protocol outlined in Choi et al. 2018^59^. Cydippids at different stages of development were fixed using the same procedure as for the colorimetric *in situ* hybridization and then stored in 100 % methanol at –20 °C until further processing. For rehydration, specimens were sequentially rinsed for 5 minutes each in 70 % methanol/30 % PTw, then 30% methanol/70% PTw, followed by three washes in PTw alone. Once rehydrated, cydippids were washed three times for 5 minutes in 5× SSCT buffer (saline-sodium citrate with 0.1% Tween-20). Prehybridization was conducted by incubating samples in HCR Probe Hybridization Buffer (Molecular Instruments, Inc., USA) for 30 minutes at 37 °C. Cydippids were then transferred into fresh HCR Probe Hybridization Buffer containing the DNA probe set (0.8 pmol per 100 µL, Molecular Instruments, Inc., USA) and allowed to hybridize overnight at 37 °C. The next day, cydippids were washed three times for 10 minutes and twice for 30 minutes in Probe Wash Buffer (Molecular Instruments, Inc., USA) at 37 °C, followed by five 5-minute washes in 5× SSCT at room temperature. Samples were then incubated for 30 minutes in Signal Amplification Buffer (Molecular Instruments, Inc., USA). Hairpin amplifiers h1 and h2 (12 pmol each, Molecular Instruments, Inc., USA) were heat-denatured at 95 °C for 90 seconds, rapidly cooled to room temperature in the dark, and subsequently added to 100 µL of Signal Amplification Buffer. Cydippids were transferred into this hairpin-containing buffer and incubated overnight at room temperature to allow amplification. Post-amplification, the cydippids were washed in 5× SSCT three times for 5 minutes and twice for 30 minutes. Finally, they were counterstained with 1 µg/mL Hoechst 33342 (TA9H97BAECD2, Targetmol Chemicals Inc. USA) and mounted in Vectashield (Vector Laboratories, Inc., USA) for imaging with a Carl Zeiss LSM 980 confocal microscope (Carl Zeiss Microscopy GmbH, Jena, Germany).

### Transmission electron microscopy

For electron microscopy, specimens were fixed overnight at 4 °C in a fixative solution containing 2.5% glutaraldehyde in 1M cacodylate buffer (pH 7.2–7.4) supplemented with 17 mg/mL NaCl and 0.025% MgCl₂. Following fixation, samples were post-fixed in 1% osmium tetroxide prepared in mQ water for 1 hour, then rinsed several times with mQ water. For transmission electron microscopy (TEM), specimens were dehydrated through a graded ethanol and isopropanol series and embedded in SPURR resin (Electron Microscopy Sciences, USA). Ultrathin sections were cut using a Leica UC7 ultramicrotome (Leica Microsystems GmbH, Wetzlar, Germany), stained 3 min with UranyLess EM stain (Electron Microscopy Sciences, USA) and 3 min 3% Lead Citrate (Electron Microscopy Sciences, USA), and examined using Zeiss-EM 900 transmission electron microscope (Electron Microscopy Center, Jena University Hospital).

### On-Grid Immunogold Labeling with Silver Enhancement

We optimised an available on-grid immunogold labelling protocols with silver enhancement^29,60,61^ for *M. leidyi*. For immunogold labelling, adult comb plates and cydippids were isolated and fixed overnight in a solution containing 4% paraformaldehyde, 0.2% glutaraldehyde, 0.4 M sucrose, and 0.1 M cacodylate buffer (pH 7.4) without agitation. The specimens were then rinsed multiple times with 0.1 M cacodylate buffer (pH 7.4). Following fixation, samples were dehydrated through a graded ethanol series and embedded in LR White resin (Electron Microscopy Sciences, Hatfield, PA, USA) at 35°C overnight. Ultrathin sections, 70 nm thick, were prepared at the Leica UC7 ultramicrotome (Leica Microsystems GmbH, Wetzlar, Germany) and placed on nickel grids coated with formvar film. Grids were first incubated with 0.05 M glycine in TBST buffer (20 mM Tris-HCl, 150 mM NaCl, 0.05% Tween-20; pH 7.9)^60^ for 10–20 minutes to quench residual aldehydes. The grids were then rinsed in a TBST buffer solution for 10 minutes and incubated overnight at 4°C with a custom-made *M. leidyi* anti-Ml-Cilin-C antibody diluted to a concentration of 1:25. Subsequent washes were performed in TBST buffer, 5×5 minutes, followed by a 1-hour incubation with Ultra Small Gold-conjugated secondary antibodies (1:50) (AURION Immuno Gold Reagents & Accessories, Wageningen, Netherlands). Grids were washed in TBST buffer, 2×5 minutes, and filtered distilled water, 2×5 minutes. Silver enhancement was carried out using the AURION R-GENT SE-EM kit (AURION Immuno Gold Reagents & Accessories, Wageningen, Netherlands) for approximately 50 minutes in light-protected conditions. After washing in filtered distilled water, 3×3 minutes, grids were air-dried and subsequently oven-dried at 40°C. Counterstaining of the ultrathin sections was performed as described above for TEM.

## Acknowledgements

This study was supported by HFSP Research Grant (RGP0041/2022, to A.H. and C.-P.H.) and DFG Grant (Nr. 519107654, to A.H.). We are grateful to Prof. Kazuo Inaba for providing us with the anti-CTENO64 antibodies. We would like to thank Prof. Dr. Elisabeth Liebler-Tenorio (FLI, Jena) and Dr. Sandor Nietzsche (Jena University Hospital) for their help with the transmission electron microscopy.

## Author contributions

A.H. – conceptualisation and funding acquisition; S.K. designed the study, performed experiments, data analysis, establishment of *Mnemiopsis leidyi* laboratory culture; S.K. wrote the first version of the manuscript; S.K. performed immunostainings and phylogenetic analysis; N.R.-K. and S.K. performed electron microscopy; T.L. performed western blot; S.K., A.K., C-P.H. performed zebrafish experiments; S.K. and A.K. performed analysis of single-cell RNA expression profiles; G.M. and L.H. produced anti-Ml-Cilin-N antibodies; All authors interpreted the results and edited the manuscript.

## Materials & Correspondence

Correspondence and requests for materials should be addressed to Prof. Dr. Andreas Hejnol or Dr. Stanislav Kremnyov.

