## Supplementary for "The ancient metazoan cytoplasmic intermediate filament protein, Cilin, shapes cilia arrangement and tissue architecture"

### Supplementary Figures

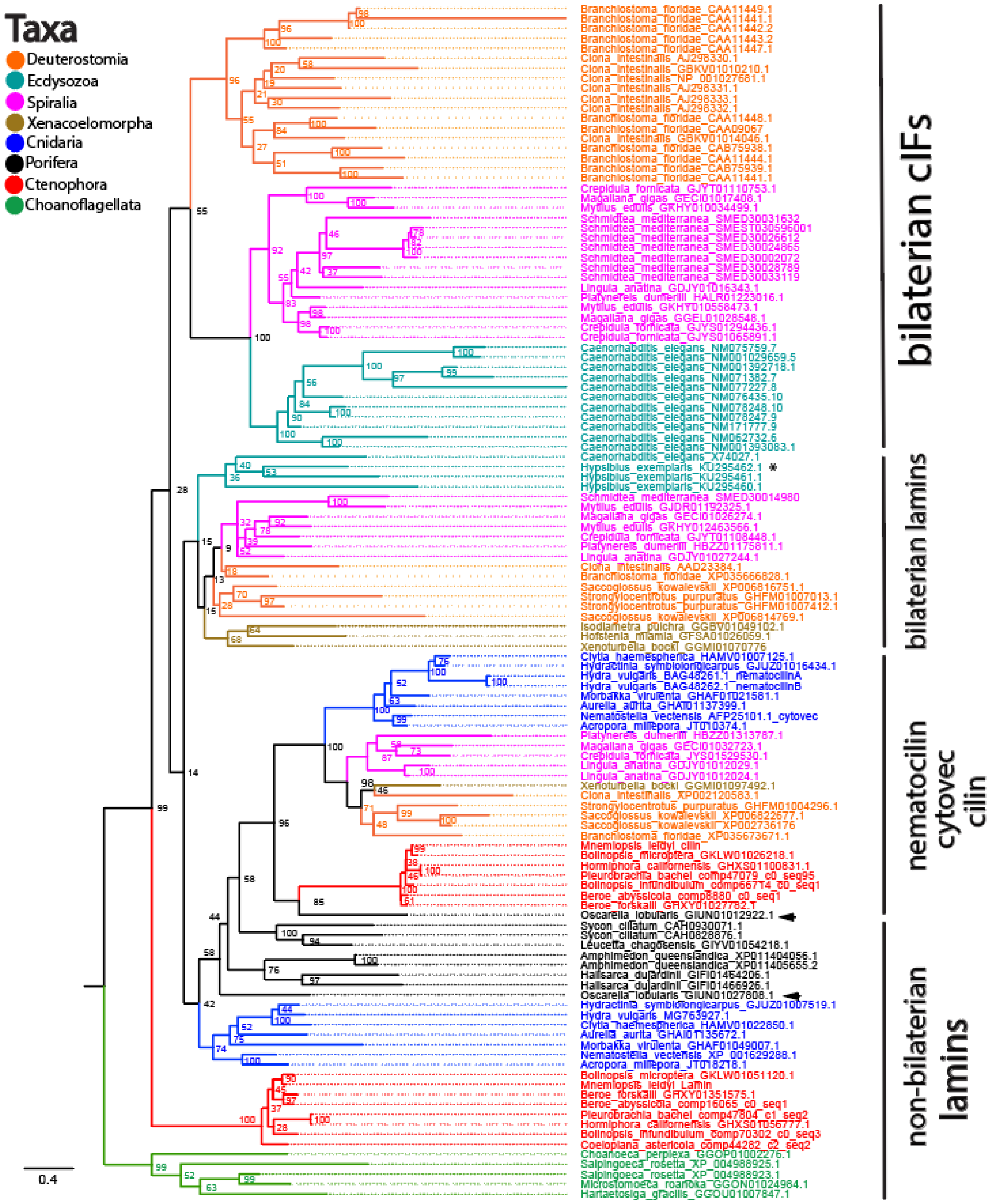


**Supp. figure 1. Phylogenetic analysis of IF proteins across Metazoa.** Maximum likelihood phylogenetic tree of IF proteins based on the rod domain, showing relationships among major metazoan lineages. Ctenophore cIF (cilin) sequences cluster with the nematocilin/cytovec group. Arrowheads indicate *Oscarella lobularis* sequences. Asterisks mark the re-evolved cIF, cytotardin, in tardigrades. Numbers at nodes are bootstrap values, shown as percentages. The scale indicates the expected number of amino acid substitutions per site.

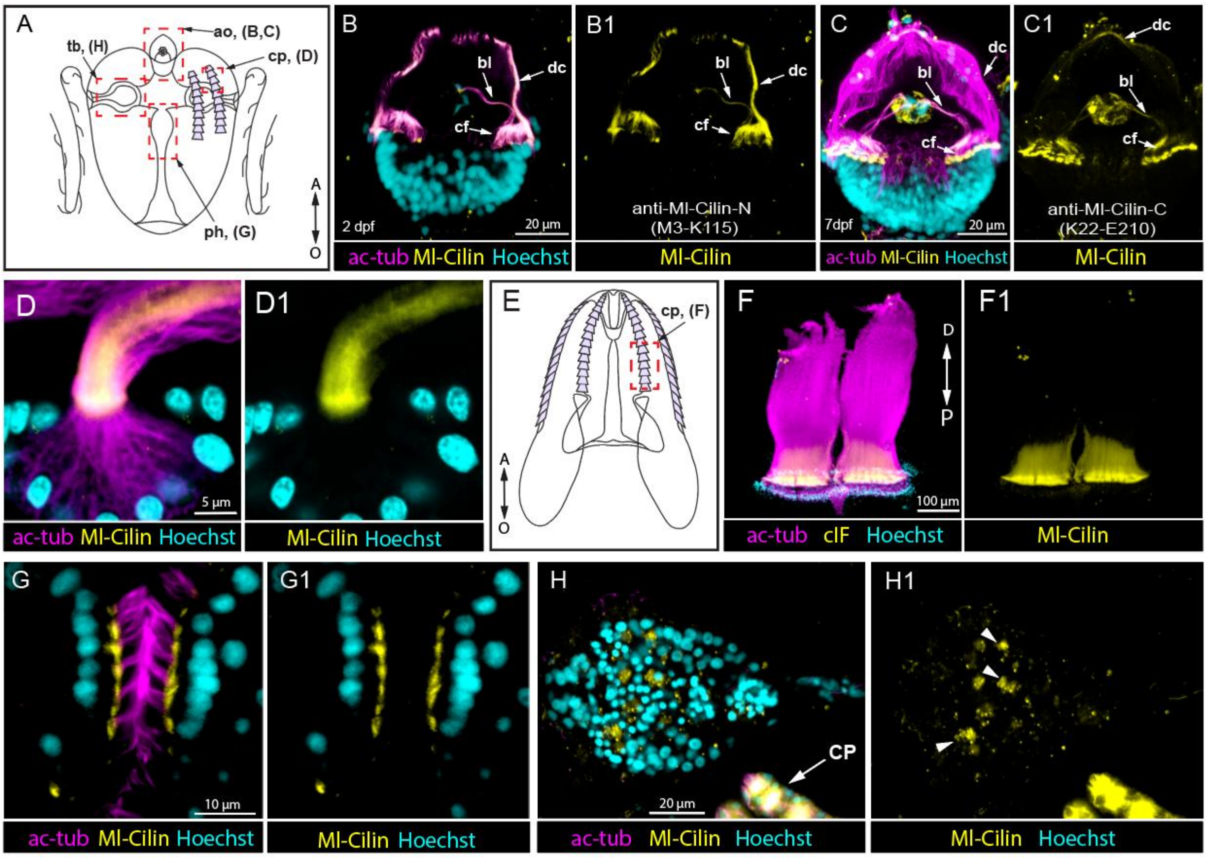


**Supp. figure 2. Immunolocalization of Ml-Cilin in 2dph cydippids using the complementary antibody anti-Ml-Cilin-N (B, B1, D-H1) and in 7 dph cydippids using anti-Ml-Cilin-C (C-C1), supporting the reliability of the main data. (A)** Schematic drawing of a *Mnemiopsis leidyi* cydippid indicating the position of the aboral organ, comb plates, tentacle bulb, and pharynx. **(B, B1)** Immunochemical staining of the aboral organ at the 2–3 dpf cydippid stage shows strong Ml-Cilin signal in the dome cilia, ciliated furrow, and balancer cilia using the anti-Ml-Cilin-N antibody. **(C, C1)** At later stages (7 dph), anti-Ml-Cilin-C also detects clF in the dome cilia, though with lower intensity compared to anti-Ml-Cilin-N, suggesting stage-specific expression or epitope accessibility. **(D, D1)** Localization of Ml-Cilin in cydippid comb plates shows signal in the proximal regions of the cilia. **(E)** Diagram of an adult *Mnemiopsis leidyi* (lobate stage) showing the location of the comb plates. **(F, F1)** Immunochemical double staining of comb plates in lobate-stage *Mnemiopsis leidyi* using antibodies against acetylated α-tubulin and Ml-Cilin. **(G, G1)** Immunochemical double staining of the pharyngeal ciliary mill reveals Ml-Cilin localized in the cytoplasm beneath the cilia. **(H, H1)** Immunochemical staining of the tentacle bulb showing Ml-Cilin expression. In **(H1)**, arrowheads indicate muscle cell precursors positive for Ml-Cilin. A – aboral; ac-tub – acetylated α-tubulin; ao – aboral organ; bc – balancer cilia; cf – ciliated furrow; cp – comb plates; dc – dome cilia; g – gut; n – nucleus; O – oral; ph – pharynx; tb – tentacle bulb.

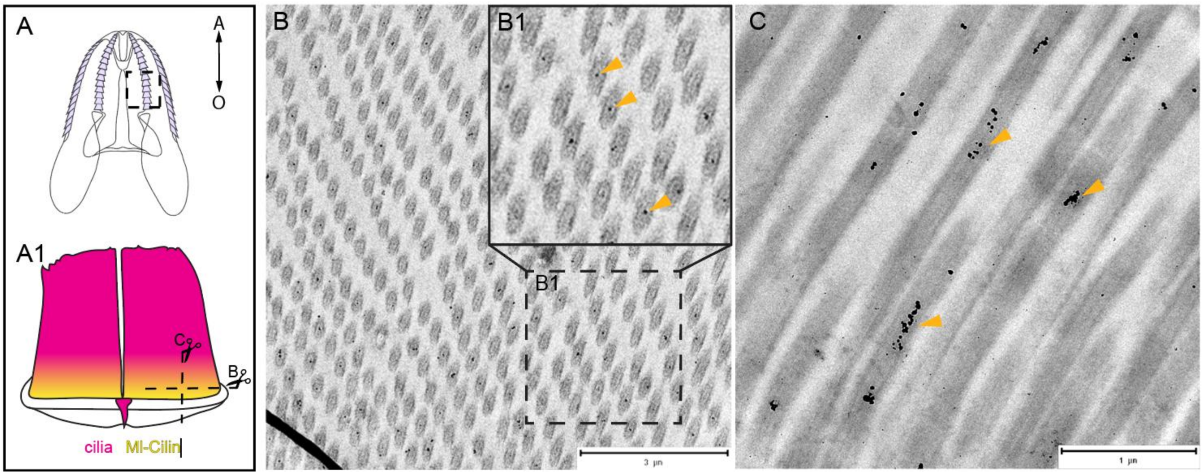


**Supp. figure 3. Immunogold labeling of Ml-Cilin in comb plate cilia. (A)** Schematic drawing of a *Mnemiopsis leidyi* cydippid indicating the position of the comb plate, and **(A1)** the level and plane of sectioning. **(B–B1)** Transverse section of the proximal region of the comb plate. **(C)** Longitudinal section of the proximal region of the comb plate. Gold particles (orange arrowheads) mark the localization of Ml-Cilin within the central region of the axoneme, supporting its association with internal ciliary structures. Labeling was performed using the anti-Ml-Cilin-C antibody.

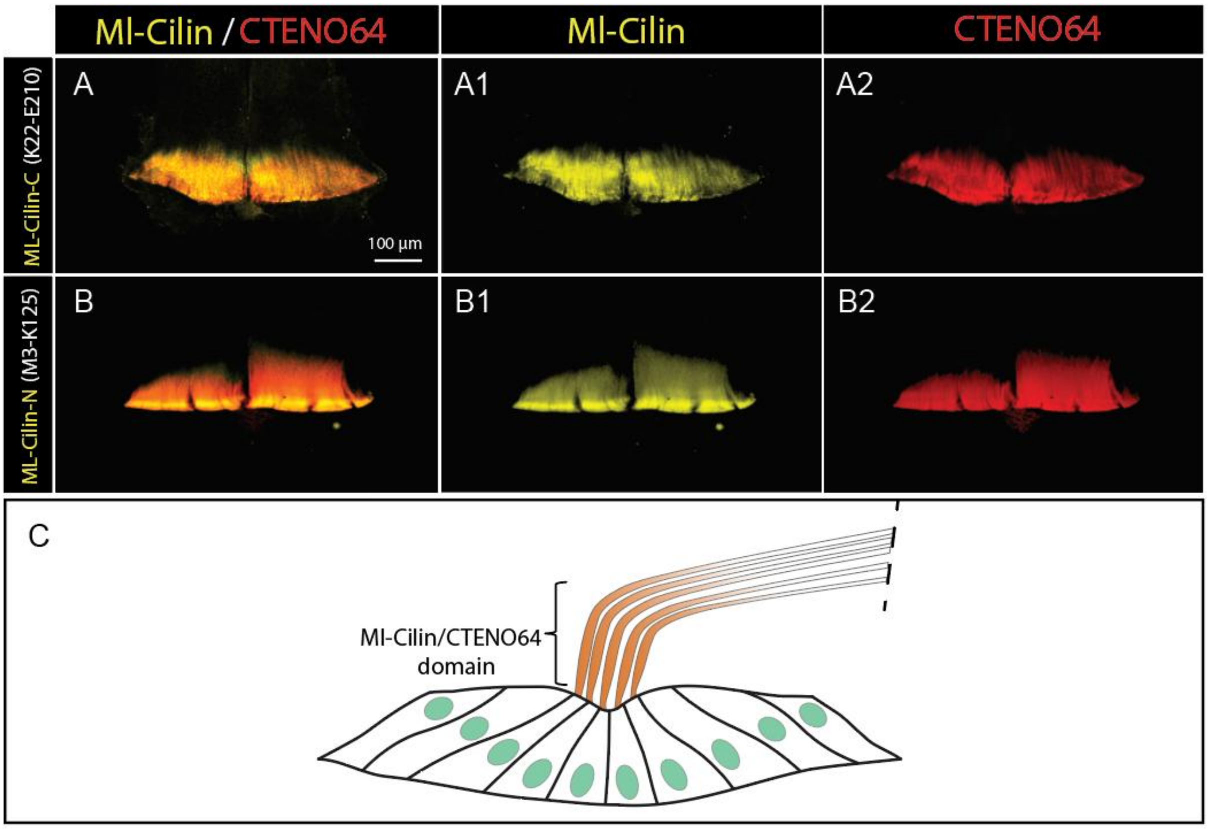


**Supp. figure 4. Ml-Cilin co-localizes with CTENO64 in comb plate cilia. (A–B2)** Double immunochemical staining shows that Ml-Cilin and CTENO64 occupy overlapping domains in the proximal region of comb plate cilia. **(C)** Schematic representation of Ml-Cilin and CTENO64 localization within the comb plate cilia. This spatial overlap in a mechanically loaded region suggests a structural reinforcement role for Ml-Cilin and CTENO64 in maintaining comb plate integrity. Staining was performed using anti-Ml-Cilin-C anti-Ml-Cilin-N, and anti-CTENO64 antibodies.

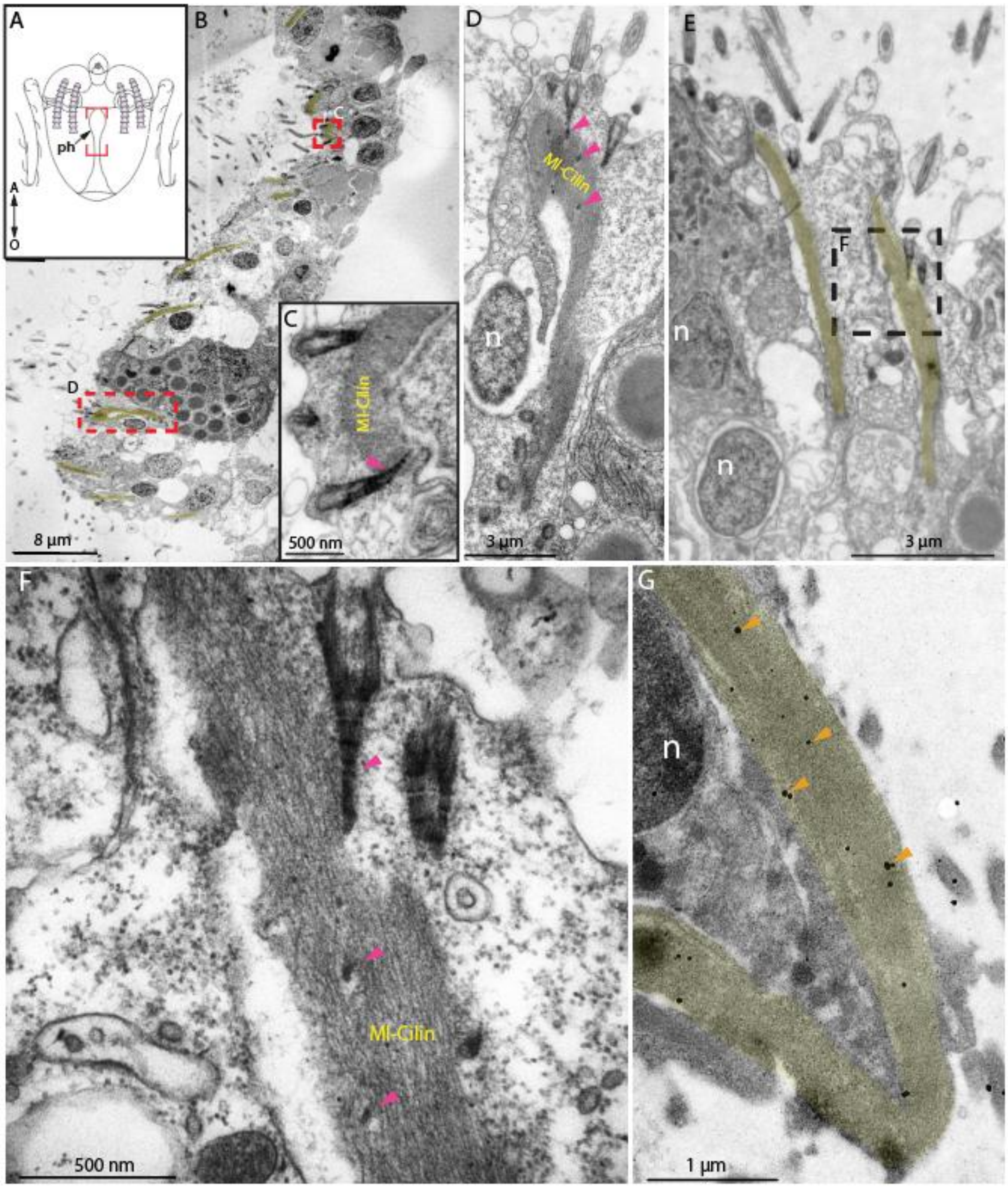


**Supp. figure 5. Ultrastructural organization of Ml-Cilin bundles in the pharyngeal ciliary mill. (A)** Schematic drawing of a *Mnemiopsis leidyi* cydippid indicating the position of the pharynx. **(B–F)** Transmission electron microscopy images show spindle-shaped bundles of filaments beneath the cilia in pharyngeal epithelial cells, aligned along cell apical–basal axis. Ciliary rootlets (purple arrowheads) are embedded within these structures. **(G)** Immunogold labeling confirms that these filamentous bundles are composed of Ml-Cilin, as indicated by gold particles (orange arrowheads) specifically associated with the bundled filaments. n – nucleus. Labeling was performed using the anti-Ml-Cilin-C antibody. A – aboral; n – nucleus; O – oral; ph – pharynx.

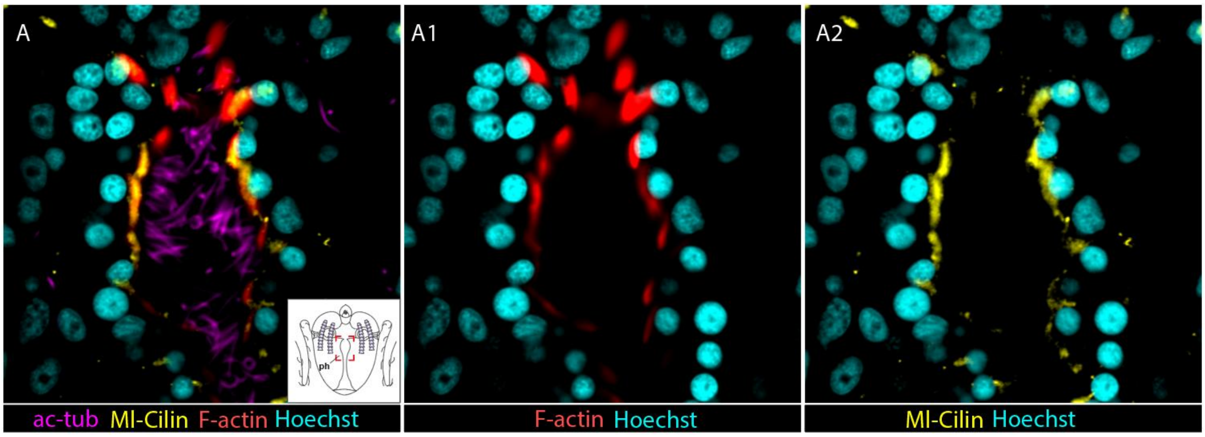


**Supp. figure 6. Co-localization of Ml-Cilin and F-actin in pharyngeal cells of *Mnemiopsis leidyi*. (A-A2)** Double immunochemical staining reveals overlapping distribution of Ml-Cilin and F-actin in the pharyngeal ciliary mill region. Ml-Cilin is localized to filamentous bundles beneath the cilia, partially overlapping with F-actin staining. This co-localization suggests that the fibrous structures observed are composite bundles composed of both cytoskeletal components. Staining was performed using anti-Ml-Cilin-C and phalloidin. ac-tub – acetylated α-tubulin; ph – pharynx.

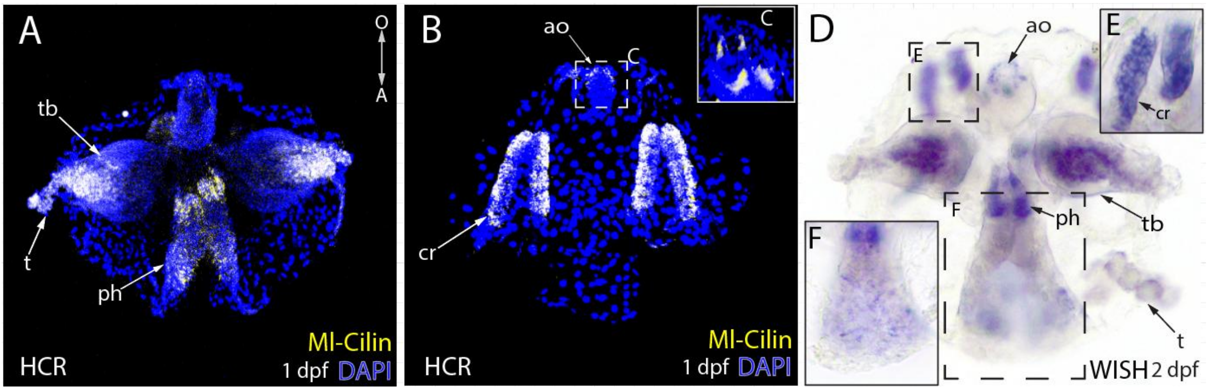


**Supp. figure 7. Gene expression pattern of Ml-Cilin in *Mnemiopsis leidyi* cydippids. (A–F)** In situ hybridization using both hybridization chain reaction **(A, B)** and colorimetric whole-mount **(D)** approaches reveals strong expression of the cIF gene in the comb rows, apical organ, ciliary mill region of the pharynx, and tentacle bulbs. The expression domains correspond to regions identified by antibody staining, supporting the reliability of the protein localization data. ao – apical organ; cr – comb rows; ph – pharynx; t – tentacle; tb – tentacle bulbs.

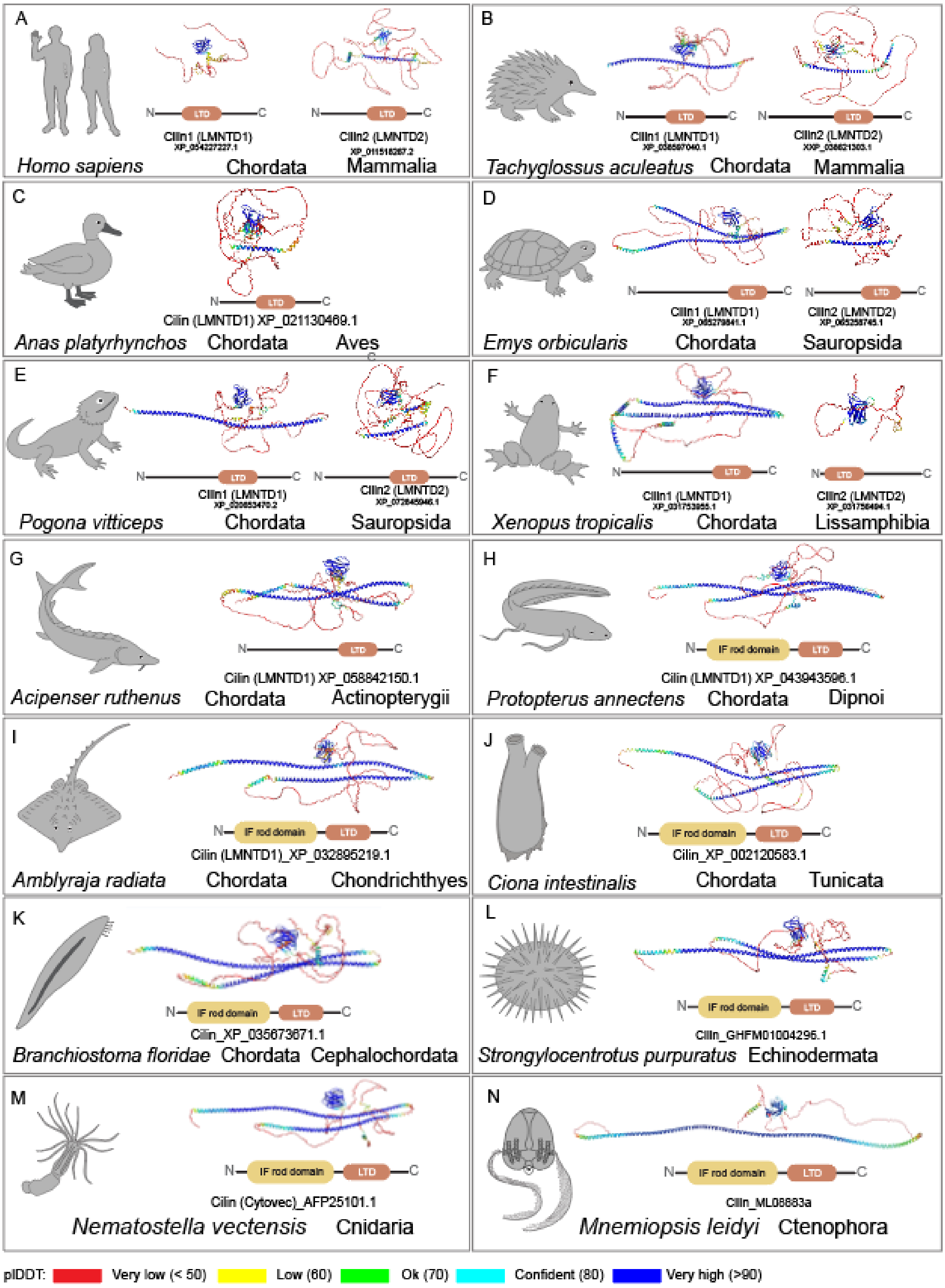


**Supp. figure 8. Predicted 3D structures of cilins and LMNTD proteins generated using AlphaFold2. (A–I)** Structural models of vertebrate LMNTD1 and LMNTD2 proteins. **(J,K)** Predicted structures of cytoplasmic intermediate filament proteins from **(J)** *Ciona intestinalis* and **(K)** Branchiostoma *floridae*, based on sequences that fall within the nematocilin/cytovec/cilin clade. **(M, N)** Predicted structures of cytoplasmic intermediate filament proteins of **(M)** *Nematostella vectensis* and **(N)** *Mnemiopsis leidyi.* We propose a new nomenclature for these proteins; for details, see the Discussion section. The schematic representation of domain structures is based on data of InterPro and the NCBI Conserved Domain Database. Notably, although LMNTD1 proteins of *Tachyglossus aculeatus*, *Emys orbicularis*, *Pogona vitticeps*, *Xenopus tropicalis* and *Acipenser ruthenus* contain α-helix segments typical for intermediate filament rod domains, these regions are not classified as filament domains by either InterPro or the NCBI Conserved Domain Database.

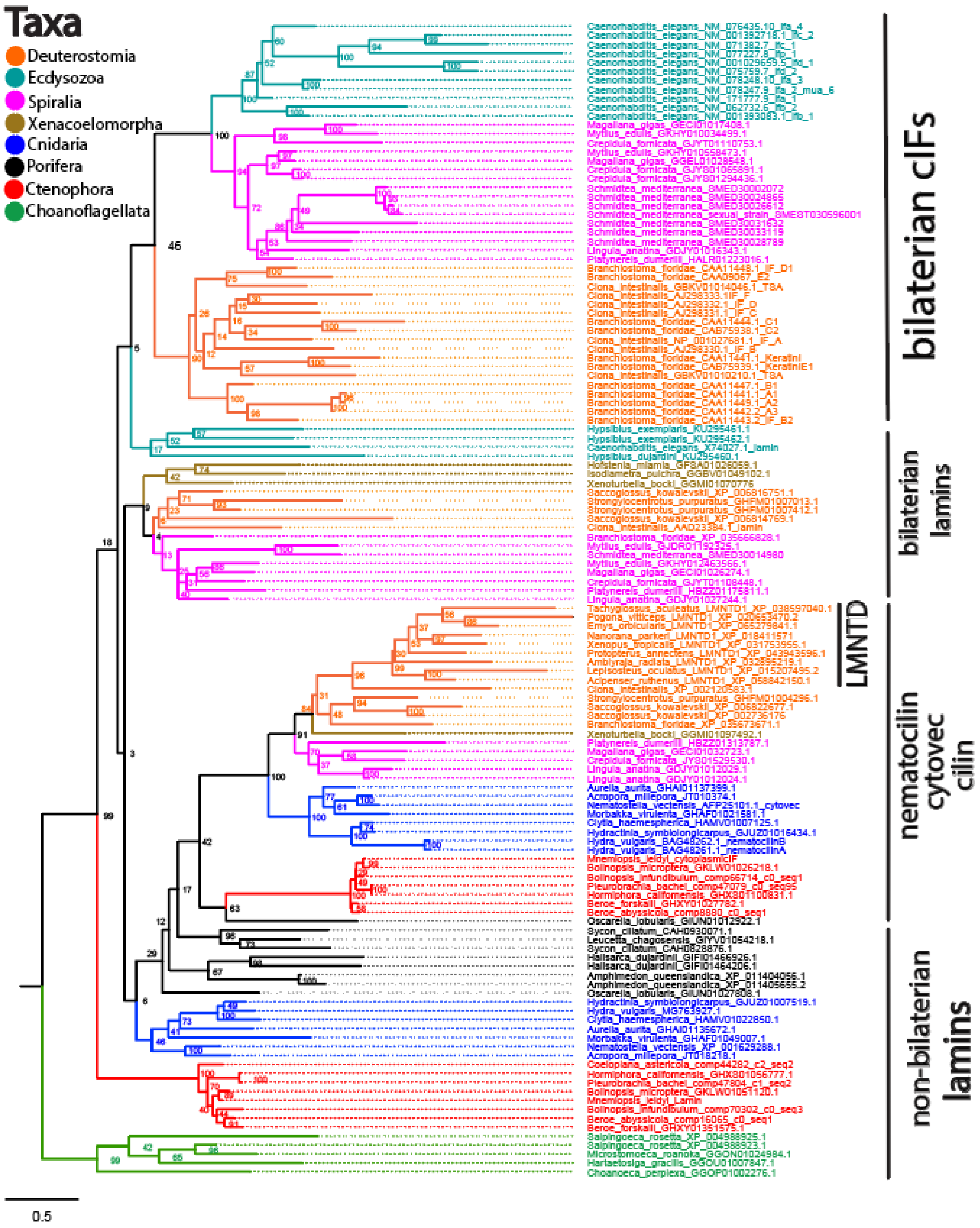


**Supp. figure 9. Phylogenetic analysis of LMNTD and related cIF proteins.** Maximum likelihood phylogenetic tree based on the N-terminal region of proteins containing an α-helical segment. All LMNTD sequences, form a monophyletic group within the nematocilin/cytovec/cilin clade, sister to *Ciona intestinalis* (bootstrap = 96) supporting their homology and shared evolutionary origin.

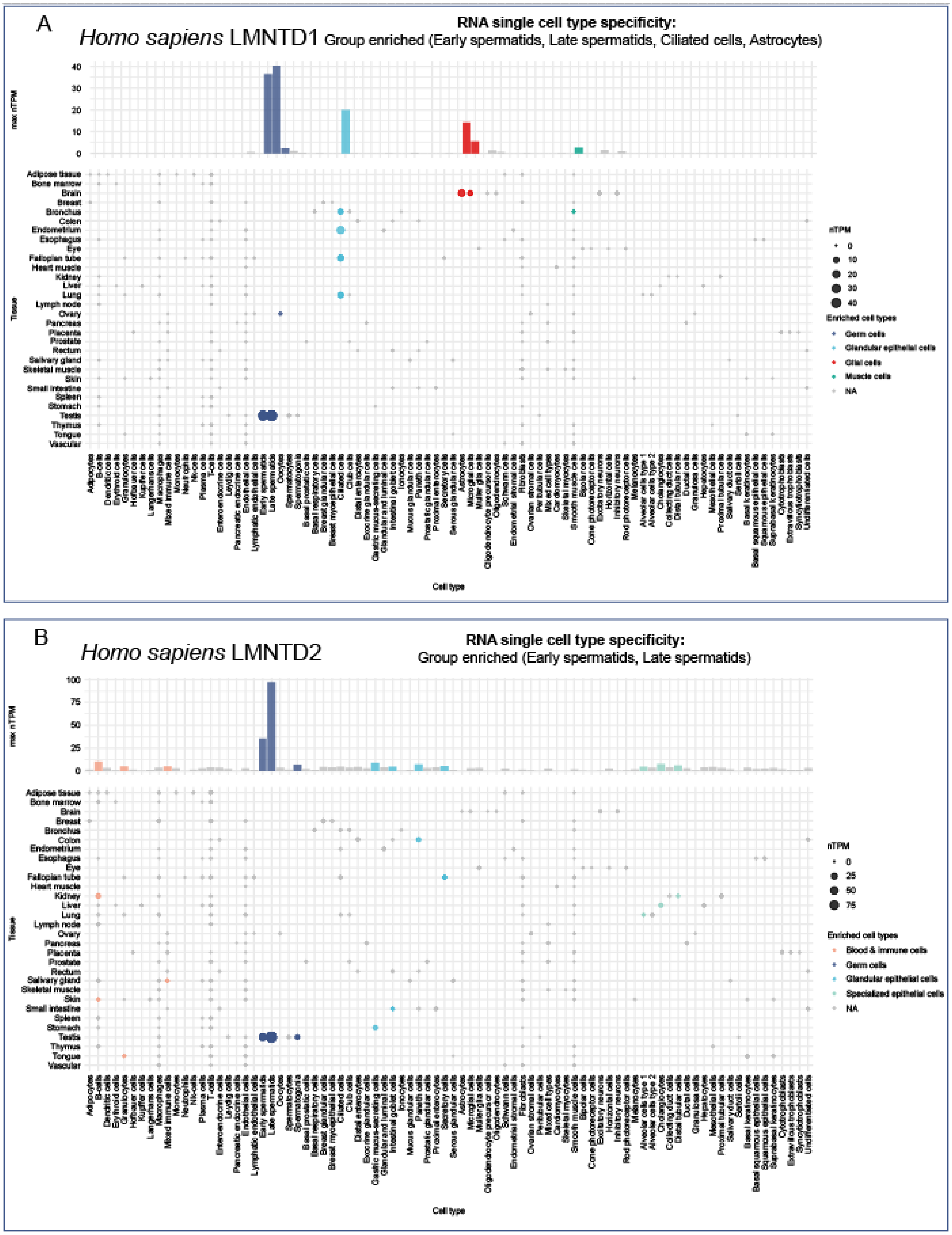


**Supp. Figure 10. Single-cell RNA expression profiles of human LMNTD1 and LMNTD2 proteins. (A)** LMNTD1 expression is enriched in early and late spermatids, as well as in multiple types of ciliated epithelial cells, including those of the endometrium, fallopian tubes, lungs, and bronchus. Additional expression is observed in astrocytes. **(B)** LMNTD2 expression is restricted primarily to early and late spermatids**.**

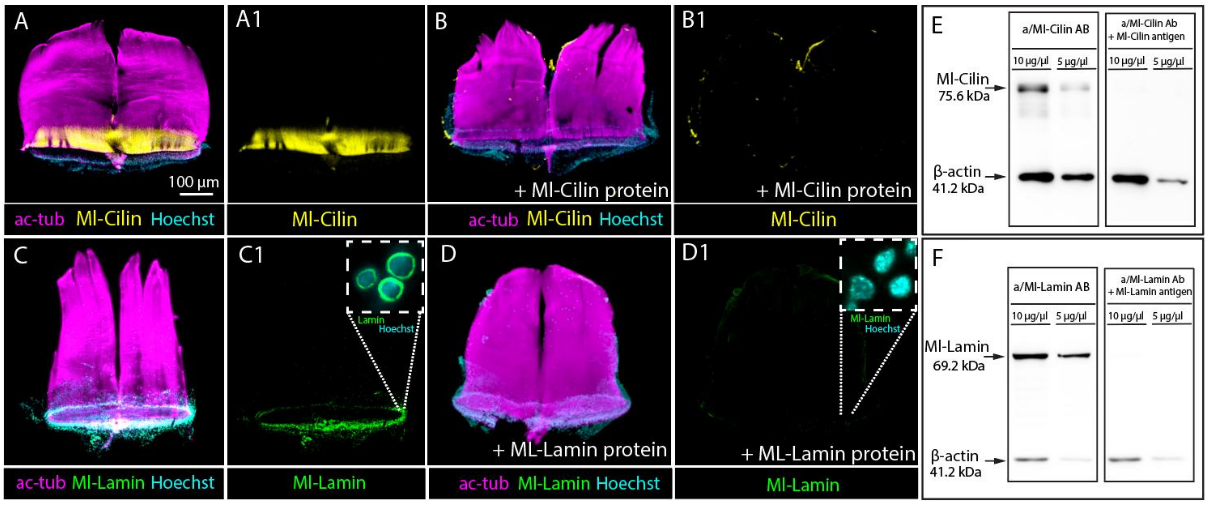


**Supp. figure 11. Validation of antibody specificity by preabsorption control and western blot. (A, A1, C, C1)** Immunochemical staining of adult comb plates using **(A, A1)** anti-Ml-Cilin-C and **(C, C1)** anti-Ml-Lamin antibodies. **(B, B1, D, D1)** Preabsorption of the same antibodies with their corresponding immunogen peptides results in complete loss of target signal, confirming staining specificity. **(E, F)** Western blot of **(E)** anti-Ml-Cilin-C and **(F)** anti-Ml-Lamin antibodies. ac-tub – acetylated α-tubulin.
